# Compressive Temporal Summation in Human Visual Cortex

**DOI:** 10.1101/157628

**Authors:** Jingyang Zhou, Noah C. Benson, Kendrick Kay, Jonathan Winawer

## Abstract

Combining sensory inputs over space and time is fundamental to vision. Population receptive field models have been successful in characterizing spatial encoding throughout the human visual pathways. A parallel question—how visual areas in the human brain process information distributed over time—has received less attention. One challenge is that the most widely used neuroimaging method—fMRI—has coarse temporal resolution compared to the time-scale of neural dynamics. Here, via carefully controlled temporally modulated stimuli, we show that information about temporal processing can be readily derived from fMRI signal amplitudes in male and female subjects. We find that all visual areas exhibit sub-additive summation, whereby responses to longer stimuli are less than the linear prediction from briefer stimuli. We also find fMRI evidence that the neural response to two stimuli is reduced for brief interstimulus intervals (indicating adaptation). These effects are more pronounced in visual areas anterior to V1-V3. Finally, we develop a general model that shows how these effects can be captured with two simple operations: temporal summation followed by a compressive nonlinearity. This model operates for arbitrary temporal stimulation patterns and provides a simple and interpretable set of computations that can be used to characterize neural response properties across the visual hierarchy. Importantly, compressive temporal summation directly parallels earlier findings of compressive spatial summation in visual cortex describing responses to stimuli distributed across space. This indicates that for space and time, cortex uses a similar processing strategy to achieve higher-level and increasingly invariant representations of the visual world.

**Significance statement:** Combining sensory inputs over time is fundamental to seeing. Two important temporal phenomena are *summation*, the accumulation of sensory inputs over time, and *adaptation*, a response reduction for repeated or sustained stimuli. We investigated these phenomena in the human visual system using fMRI. We built predictive models that operate on arbitrary temporal patterns of stimulation using two simple computations: temporal summation followed by a compressive nonlinearity. Our new temporal compressive summation model captures (1) subadditive temporal summation, and (2) adaptation. We show that the model accounts for systematic differences in these phenomena across visual areas. Finally, we show that for space and time, the visual system uses a similar strategy to achieve increasingly invariant representations of the visual world.

## 1 Introduction

A fundamental task of the visual system is to combine sensory information distributed across space and time. How neural responses sum inputs across space has been well characterized, with several robust phenomena. First, spatial summation in visual cortex is subadditive: the response to two stimuli presented in different locations at the same time is less than the sum of the responses to the stimuli presented separately. This phenomenon is observed in all cortical areas studied and has been measured with both fMRI (Kastner et al., 2001; Kay et al., 2013a) and electrophysiology (Rolls and Tovee, 1995; Britten and Heuer, 1999; Heuer and Britten, 2002; Winawer et al., 2013); such nonlinearities may reflect an adaptation to achieve efficient encoding of natural images (Schwartz and Simoncelli, 2001). In addition, in higher visual areas, receptive field size increases (Maunsell and Newsome, 1987) and sub-additive summation becomes more pronounced (Kay et al., 2013a; Kay et al., 2013b). As the subadditivity becomes more pronounced in later areas and receptive fields get larger, a stimulus that occupies only a small fraction of a neural receptive field can produce a large response. As a result, responses in higher visual areas become increasingly insensitive to changes in the size and position of stimuli (Tovee et al., 1994; Grill-Spector et al., 2001; Kay et al., 2013a). The tendency towards increasing tolerance for size and position in higher areas trades off with the increasing specificity of tuning to higher level stimulus information (Rust and Dicarlo, 2010, 2012).

Here, we hypothesize that the same organizational principles for the visual cortex apply in the temporal domain (Figure 1). Just as natural images tend to vary slowly over space, image sequences typically vary slowly over time (Dong and Atick, 1995; Weiss and Adelson, 1998). As a result, an efficient code would prioritize abrupt changes in time over sustained or repeated stimuli (Snow et al., 2016); this would result in sub-additive temporal summation for sustained or repeated stimuli (also referred to as adaptation or repetition suppression). Evidence for such temporal non-linearities are abundant in single cell recordings of primary visual cortex (for example, Tolhurst et al., 1980), but have not been systemically characterized across visual areas or with a forward model. At longer time scales, the fMRI BOLD signal sums contrast patterns close to, but slightly less than, linearly (Boynton et al., 1996; Boynton et al., 2012). We hypothesize that (1) at the time scale of neuronal dynamics in sensory cortex (tens to hundreds of ms), temporal summation will be substantially subadditive, and (2) that more anterior visual areas will show greater subadditivity. This greater subadditivity in later areas will make these responses less sensitive to the precise duration and timing of a stimulus, paralleling size and position tolerance in the spatial domain. This prediction is consistent with the logic that later visual areas trade off position and duration specificity for increased tuning for high level stimulus properties.

**Figure 1.**
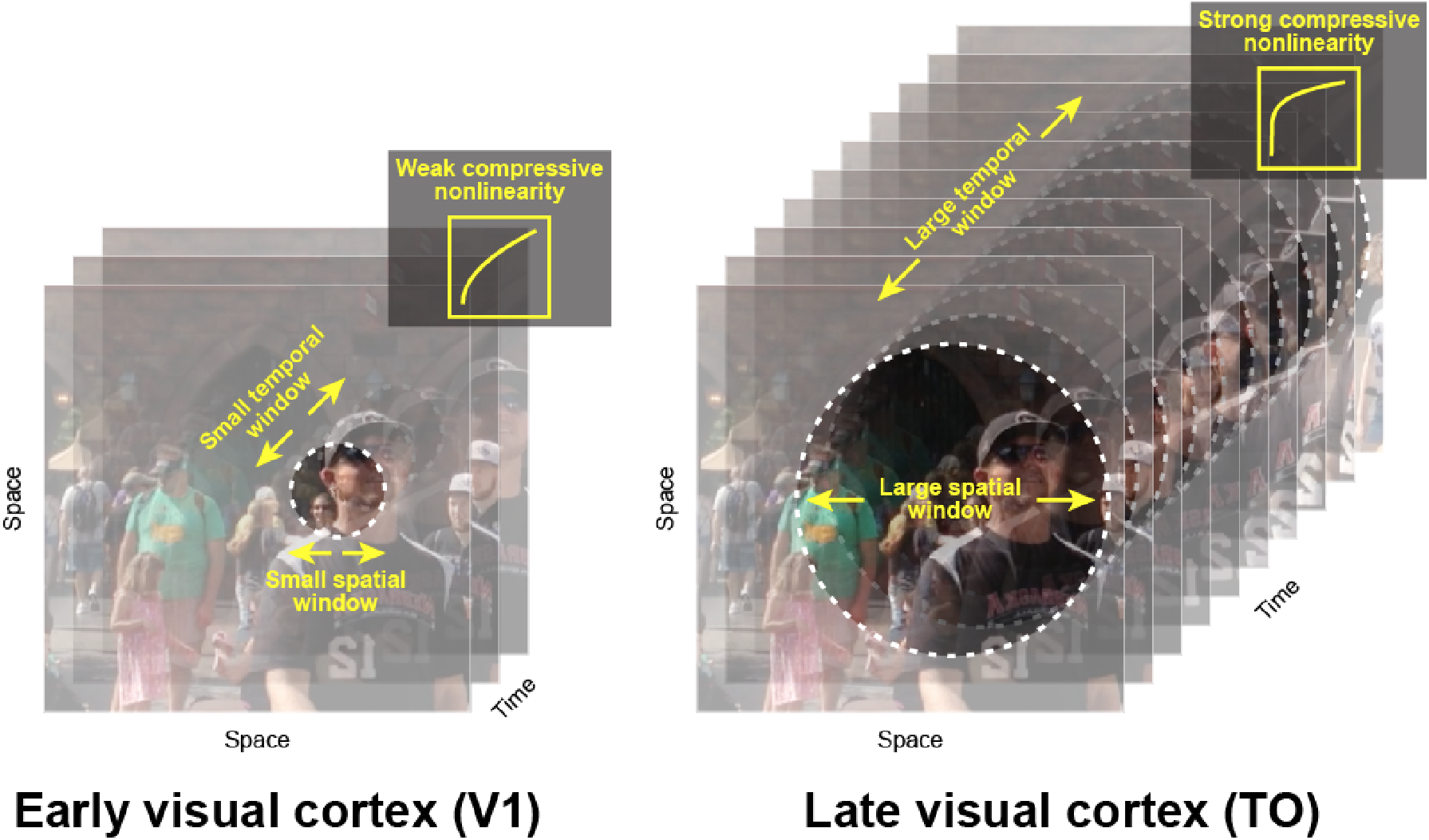
Parallels between spatial and temporal processing. It is well established that spatial receptive fields are small in V1 (left) and grow larger in later visual areas such as the temporal occipital maps (‘TO’, right). It was recently shown that there is also a gradient of an increasingly pronounced compressive summation over space from early to later areas (Kay et al., 2013a). Here, we hypothesize that temporal summation, as well as the temporal receptive field size, follows a similar pattern, with increasingly long temporal windows and more compressive summation over time in the more anterior visual areas. We propose that the combination of larger spatiotemporal windows and more compressive nonlinearities is part of a coding strategy whereby higher visual areas achieve increasing invariance to changes in stimulus size, position, and duration.

In this paper, we used fMRI to study temporal summation and adaptation. We characterized responses to brief stimuli (tens to hundreds of ms) in many visual areas, measured with fMRI, which has the advantage of being non-invasive and recording from many visual areas in parallel. To quantify and understand how temporal information is encoded across visual cortex, we implemented a temporal population receptive field (“pRF”) model which predicts the fMRI response amplitude to arbitrary stimulus time courses.

## 2 Materials and methods

### 2.1 fMRI procedure

#### Participants

Data from six experienced fMRI participants (two males and four females, age range 21-48, mean age 31) were collected at the Center for Brain Imaging (CBI) at NYU. All participants had normal or corrected-to-normal visual acuity. The experimental protocol was approved by the University Committee on Activities Involving Human Subjects, and informed written consents were obtained from all participants prior to the study. For each participant, we conducted a 1-hour session for visual field map identification and high-resolution anatomical volumes, and either one or two 1.5-hour sessions to study temporal summation. Two of the six participants (one male, one female) were included in both the main temporal summation experiment and the self-replication experiment (hence two 1.5-hour sessions). The other four participants (one male and three females) were included in only the main temporal summation experiment or the self-replication experiment.

#### Visual Stimuli

##### Stimuli

For the main experiment, stimuli were large field (24° diameter) band-pass noise patterns (centered at 3 cycles per degree), independently generated for each trial. The pattern was chosen because it was previously shown to be effective in eliciting responses in most visual areas (Kay et al., 2013b). (See Kay et al (2013b) for details on stimulus construction). A second experiment replicated all aspects of the main experiments except that the stimulus patterns differed. For this experiment, the patterns were either pink noise (1/f amplitude spectrum, random phase), or a front-view face image embedded in the pink noise background. The face stimuli were the front-facing subset of the faces used by Kay et al (2015). For both experiments, stimuli were windowed with a circular aperture (24° diameter, 768 x 768 pixels) with a raised cosine boundary (2.4 deg). All stimuli were gray scale. Stimulus generation, presentation and response recording were coded using Psychophysics Toolbox (Brainard, 1997; Pelli, 1997) and vistadisp (https://github.com/vistalab/vistadisp). We used a MacBook Air computer to control stimulus presentation and record responses from the participants (button presses) during the experiment.

##### Display

Stimuli were displayed via an LCD projector (Eiki LC_XG250; resolution: 1024 x 768 pixels; refresh rate: 60 Hz) onto a back-projection screen in the bore of the magnet. Participants, at a viewing distance of ~58 cm, viewed the screen (field of view, horizontal: ~32°, vertical: ~24°) through an angled mirror. The images were confined to a circular region with a radius of 12º. The display was calibrated and gamma corrected using a linearized lookup table.

##### Fixation task

To stabilize attention level across scans and across participants during the main experiment, all participants were instructed to do a one-back digit task at the center of fixation throughout the experiment, as in previous publications (Kay et al., 2013a; Kay et al., 2013b). The digit (0.24° x 0.24°) was presented at the center of a neutral gray disk (0.47° diameter). Within a scan, each digit (randomly selected from 0 to 9) was on for 0.5 second, off for 0.167 second before the next digit appeared at the same location. Participants were asked to press a button when a digit repeated. Digit repetition occurred around 2-3%, with no more than two identical digits being presented successively. To reduce visual adaptation, all digits alternated between black and white, and on average participants pressed a button every 30 seconds. During the retinotopy task, the fixation alternated pseudo-randomly between red and green (switches on average every 3s), and the participant pressed a button to indicate color changes.

#### Experimental Design

We used a randomized event-related experimental design (Figure 2A–B) to prevent participants from anticipating the stimulus conditions. An event is a stimulus presented according to one of thirteen distinct time courses (< 800 ms in total), either a single pulse with variable duration or a double pulse with fixed duration and variable inter-stimulus interval (ISI). Durations and ISIs were powers of 2 times the monitor dwell time (1/60 s). Each pulse in the double-pulse stimuli lasted 134 ms. The 0-ms stimulus was a blank (zero-contrast, mean luminance, and hence identical to the preceding and subsequent blank screen between stimulus events). For the main experiment, each participant completed seven scans, and within a scan, each temporal event repeated 4 times. A temporal event started with the onset of a pattern image, and the inter-trial interval (stimulus plus subsequent blank) was always 4.5 seconds. For stimuli with two pulses, the two noise patterns were identical. The design was identical for the self-replication experiment, except that each time course repeated three times per scan instead of 4, and each participant completed 6 scans.

**Figure 2.**
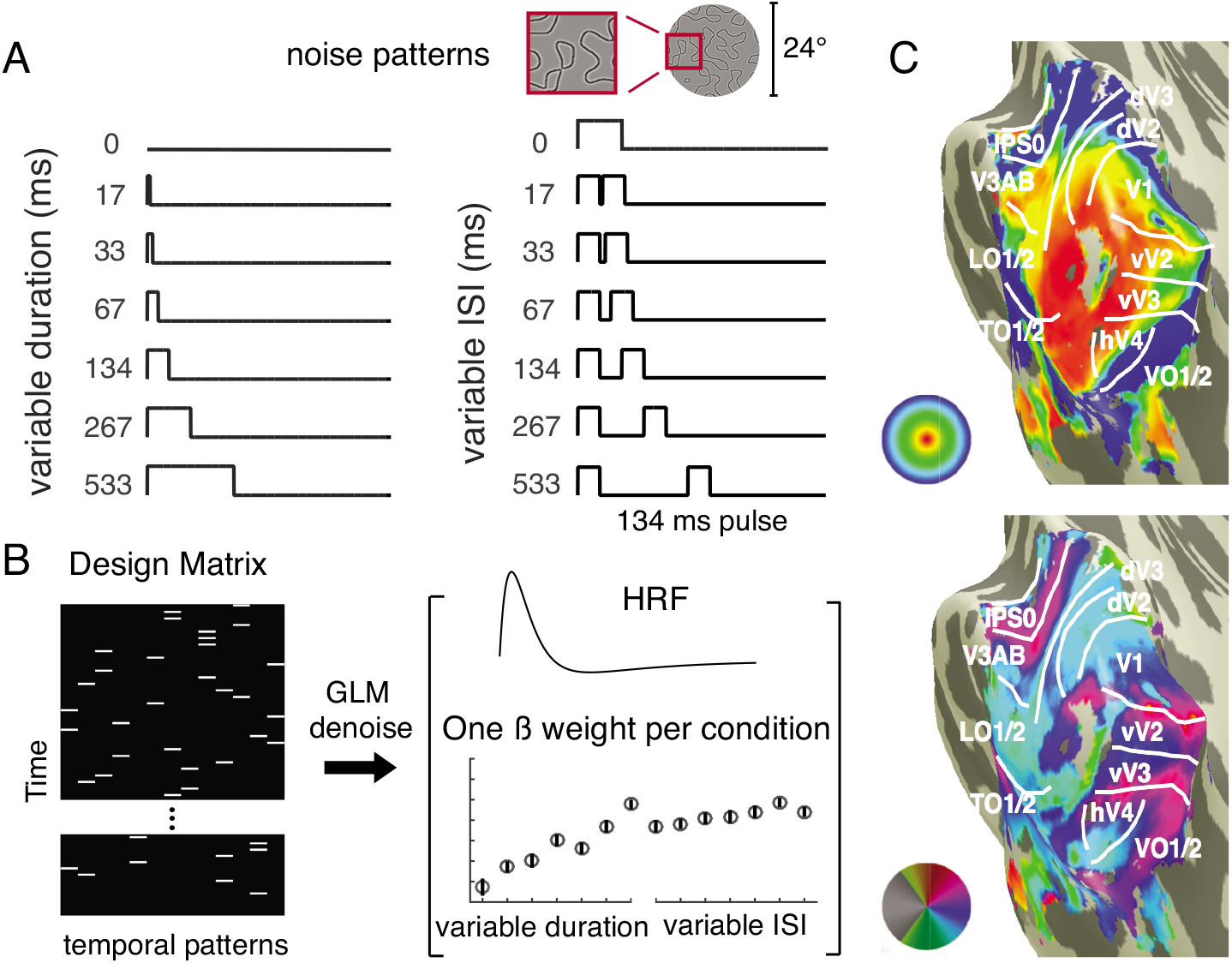
Experimental design and analysis. *(A)* Participants were presented with one or two pulses of large field (24°) spatial contrast patterns. One-pulse stimuli were of varying durations and two-pulse stimuli were of varying ISI (with each pulse lasting 134ms). *(B)* The temporal conditions were presented in random order, indicated by the white bars in the 13-column design matrix (one column per temporal condition). To analyze the data, we extracted a ß-weight for each temporal condition per area using a variant of the general linear model, GLM denoise. *(C)* Nine visual field maps or visual field maps pairs were bilaterally identified for each participant (V1; V2; V3; hV4; VO-1/2; V3A/B; IPS-0/1; LO-1/2; TO-1/2).

#### MRI Data Acquisition

All fMRI data were acquired at NYU Center for Brain Imaging (CBI) using a Siemens Allegra 3T head-only scanner with a Nova Medical phased array, 8-channel receive surface coil (NMSC072). For each participant, we collected functional images (1500 ms TR, 30 ms TE, and 72-degree flip angle). Voxels were 2.5mm^3^ isotopic, with 24 slices. The slice prescription covered most of the occipital lobe, and the posterior part of both the temporal and parietal lobes. Images were corrected for B0 field inhomogeneity using CBI algorithms during offline image reconstruction.

In a separate session, we acquired two to three T1-weighted whole brain anatomical scans (MPRAGE sequence; 1mm^3^). Additionally, a T1-weighted “inplane” image was collected with the same slice prescription as the functional scans to aid alignment of the functional images to the high-resolution T1-weighted anatomical images. This scan had an inplane resolution of 1.25 x 1.25 mm and a slice thickness of 2.5 mm.

#### Data Preprocessing and Analysis

##### Data preprocessing

We co-registered and segmented the T1-weighted whole brain anatomical images into gray and white matter voxels using FreeSurfer’s auto-segmentation algorithm (surfer.nmr.mgh.havard.edu). Using custom software, vistasoft (https://github.com/vistalab/vistasoft), the functional data were slice-time corrected by resampling the time series in each slice to the center of each 1.5s volume. Data were then motion-corrected by co-registering all volumes of all scans to the first volume of the first scan. The first 8 volumes (12 seconds) of each scan were discarded for analysis to allow longitudinal magnetization and stabilized hemodynamic response.

##### GLM analysis

For analysis of the temporal summation functional data, we used a variant of the GLM procedure—GLM denoise (Kay et al., 2013c), a technique that improves signal-to-noise ratios by entering noise regressors into the GLM analysis. Noise regressors were selected by performing principle component analysis on voxels whose activities were unrelated to the task. The optimal number of noise regressors was selected based on cross-validated R^2^ improvement (coefficient of determination). The input to GLM denoise was the pre-processed EPI data and a design matrix for each scan (13 distinct temporal profiles x number of volumes per scan), and the output was Beta-weights for each temporal profile for each voxel, bootstrapped 100 times across scans (Figure 2B). For analysis, we normalized all 13 Beta-weights per voxel by the vector length and selected a subset of voxels (see *Voxel selection*). We then averaged the Beta-weights for a given temporal condition from the first bootstrap across voxels within each ROI and across all participants to get a mean; this gives one estimate of the mean response per ROI for a given condition. This was repeated for each condition, and then repeated for each of the 100 bootstraps, yielding a matrix of 100 x 13 for each ROI (bootstraps by temporal condition). GLM denoise was not applied to the visual field map measurements, since these experiments did not have an event-related design, and hence are not amenable to a GLM analysis.

##### ROI identification

We fitted a linear population (‘pRF’) model (Dumoulin and Wandell, 2008) to each subject’s retinotopy data (average of two scans). We made an initial guess of ROI locations by first projecting the maximum likelihood probabilistic atlas from Wang et al (2015) onto the cortical surface. Then we visualized eccentricity and polar angle maps derived from the pRF model fits and modified ROI boundaries based on visual inspection. For each participant, we defined nine bilateral ROIs (V1, V2, V3, hV4, VO-1/2, LO-1/2, TO-1/2, IPS-0/1) (Figure 2C). For the second experiment (self-replication), in addition to the nine ROIs from the main experiment, we also identified a bilateral face-selective region of interest. This ROI included face-selective voxels in the inferior occipital gyrus (‘IOG-faces, or ‘Occipital Face Area’) and in the posterior fusiform (pFus, or ‘FFA-1’) (Gauthier et al., 2000; Weiner and Grill-Spector, 2010). We identified these areas by taking the difference between the mean fMRI response to all face images and the mean response to all noise images, and then thresholding the difference for voxels at the two anatomical locations (IOG and pFus), as described previously (Weiner and Grill-Spector, 2010; Kay et al., 2015).

##### Voxel selection

All analyses were restricted to voxels that satisfy the following three criteria. First voxels must be located within 2-10° (eccentricity) based on the pRF model. Second, voxels must have a positive Beta-weight for the average across all non-blank temporal conditions (and averaged across bootstraps). The bootstraps, computed by GLM denoise, were derived by sampling with replacement from the repeated scans. Third, voxels must have > 2% GLM R^2^. Voxels that satisfy all criteria were averaged within a participant to yield 13 beta weights per ROI per participant per 100 bootstraps. The data were then averaged across participants. Averaging within a participant prior to averaging across participants ensures that the contribution for each participant has the same weight, irrespective of the numbers of voxels per participant.

#### Hemodynamic response function (HRF) for individual ROIs

In section 3.8, we estimated an HRF for each ROI to test whether the use of ROI-specific HRFs, rather than a single HRF for each subject, altered the pattern of results. We estimated the HRFs using two experiments, the retinotopic mapping experiment and the temporal summation experiment. In both cases, we assumed that the HRF was parameterized by the difference between two gamma functions with five free parameters (Friston et al., 1998; Worsley et al., 2002). For the retinotopy HRF, we used the vistasoft pRF code, which uses an iterative approach alternating between fitting the pRF parameters and the HRF parameters: first the HRF is assumed to have default parameters for all voxels and the pRF parameters are fit; then the pRF parameters are fixed and the HRF is found which maximizes the mean variance explained across voxels in an ROI; finally, the HRF parameters are fixed and the pRF parameters are refit. This procedure was done separately for each ROI.

To estimate HRFs from the temporal experiment, we first selected voxels for each ROI as described in *voxel selection*. We averaged the output time series from *GLMdenoise* (0-mean, polynomial detrended, and noise PCs regressed-out) from the selected voxels within each ROI to estimate a set of Beta-weights. As with the retinotopy experiment, we estimated the HRF and a set of Beta-weights for each ROI using an iterative procedure. Each HRF was parameterized using the difference between two gamma functions with five free parameters (same as for retinotopy), and was seeded using the same vistasoft default parameters (see *rmHrfTwogamms.m*). The iterative fitting procedure terminates when the two types of fits converge.

### 2.2 Temporal pRF Models

We used two variants of a temporal pRF model, one linear and one non-linear, to predict neuronal summation measured using fMRI. All model forms take the time course of a spatial contrast pattern as input (*T*_input_), and produce a predicted neuronal response time course as output. To predict the fMRI data (BOLD), we summed the predicted time course within a trial (< 1 s) to yield one number per temporal condition. For model fitting, these numbers were compared to the fMRI Beta-weights, derived from the GLM denoise analysis.

#### Models

##### Linear model

The linear model prediction is computed by convolving a neuronal impulse response function (IRF) with the stimulus time course (T_input_), and scaling by a gain factor (*g*)

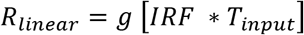

The time course is then summed for the fMRI predictions (plus an error term, *e*):

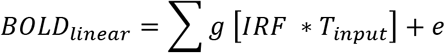

For the IRF, we assumed a gamma function, parameterized by _!_, of the form,

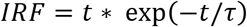

Because the IRF was assumed to have unit area, the specific shape of the IRF has no effect on the predictions of the linear model, and the prediction reduces to:

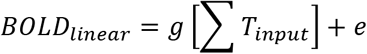

and the only value solved for is the gain factor, *g*

##### Compressive temporal summation model (CTS)

The CTS model is computed with a linear convolution, followed by a divisive normalization. The linear step is identical to the linear model. For the divisive normalization:

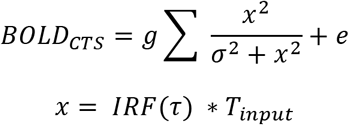

we solved for τ, σ, and gain factor *g* for the CTS model. In section 3.9, we implemented two additional variations of this model. In the first variation, we relaxed the exponent from 2 to *n*, and fitted 4 parameters, τ, σ, *n*, and the gain factor *g*.

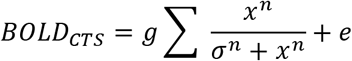

In the second variation, we allowed the exponent in the numerator to be different from that in the denominator, and we fitted 5 parameters, τ, σ, *n, m*, and *g*.

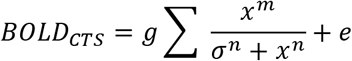

##### Compressive summation model (CTS) with power law implementation

In section 3.9, we implemented the CTS model with a power law nonlinearity rather than divisive normalization. To compute the model predicted neuronal response, we first computed the linear response by convolving an IRF (gamma function with variable time to peak τ) with an input stimulus time course, identical to the normalization implementation. Then an exponent εϵis applied point-wise to the predicted linear output.

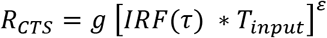

To fit the CTS model with a power law to the fMRI data, we again summed the predicted response time series:

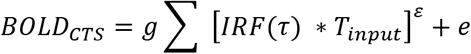

and solved for τ_1_, ε, and *g*.

Because of the nonlinearity, the specific shape of the impulse response function does matter, in contrast to the linear model. This version of the CTS model is identical to the normalization implementation, except that the shape of the compressive non-linearity due to the power law is slightly different from the shape obtained using divisive normalization.

##### Two temporal channels (TTC) model

We implemented a two-temporal-channels model, previously used for fitting V1 responses to temporal variation in luminance (Horiguchi et al., 2009). The two temporal channels model consists of a linear combination of the outputs from a sustained and a transient temporal channel. The output of the sustained channel is computed by convolving an impulse response function with the stimulus. The output of the transient channel is computed by convolving the transient impulse response function with the stimulus time course, then squared point-wise in time. The sustained IRF has a positive mean and the transient IRF has a zero mean, as implemented by (Horiguchi et al., 2009) for fMRI data. This IRF form was previously used for modeling psychophysical data (Watson, 1982; McKee and Taylor, 1984). The outputs from both channels are weighted by parameter *a* and *b*.

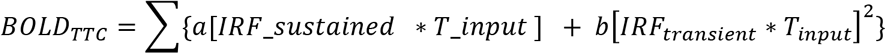

The form of both the sustained and the transient channel IRFs is the same:

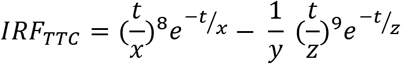

with *t* being time, and *x*, *y* and *z* take values [3.29, 14, and 3.85] respectively for the sustained IRF, and [2.75, 11, 3.18] for the transient IRF.

#### Parameter estimation and model fitting

All models except the linear model were fit in two steps, a grid fit followed by a search fit, as described below. (Also, see script *trf_fitModel.m.)*

##### CTS model – normalization implementation

For the grid fit, we computed the model response to the 13 temporal conditions for 100 combinations of τϵand σϵ(τϵvalues from 10^-3^ to 1 with 10 equal steps, and σϵfrom 10^-3^ to 0.5 with 10 equal steps). For each ROI, the parameter pair generating the predictions with the highest correlation to the data was then used as a seed for the search fit. This was repeated 100 times per ROI, once for each bootstrap of the data. See *trf_gridFit.m* and *trf_fineFit.m*.

We then did a search fit using a bounded nonlinear search algorithm in Matlab (*fminsearchbnd.m)*, 100 times per ROI, using the 100 sets of bootstrapped Beta-weights, and the 100 seed values as derived above. The search finds the parameters that minimize the squared error between predicted and measured Beta-weights. The lower bound used for the search fit is [10^-3^, 10^-4^, 0] for τ, σ, and g, a scaling factor. The upper bound was [1, 1, 1]. This gave us 100 estimates of each model parameter for each ROI, which we summarized by the median and 50% confidence interval.

Additionally, for section 3.9, we implemented the normalization model with additional free parameters. To fit the first variation of the normalization model (free parameters: τ, σ, *n*, *g*), we used the same ten steps grid fit for τϵand σϵas in the previous model. For *n*, a ten-step grid with equal steps from 0.1 and 6 was used. In the search fit stage, same bound was used for τ, σ, and *g* as in the previous model, and the bound for *n* was [0, 10]. To fit the second variation of the normalization model (free parameters: τ, σ, *n*, *m, g*), the same ten-step grid was used for both *n* and *m* in the search stage, and the same bound for both exponents was used in the search stage.

##### CTS model – exponential implementation

For the grid fit, we computed the model responses to the 13 temporal conditions for 100 combinations of τϵand εϵ(τϵvalues from 10-3 to 1 with 10 equal steps; εϵvalues from 10-3 to 2 with 10 equal steps). The procedure to select the best fitting parameter pair and the search fit step is the same as above. The lower bound we used for the search fit is [0.02, 0.001, 0] for τ, ε, and g (a scaling factor). The upper bound was [1, 2, 1]. See *trf_gridFit.m* and *trf_fineFit.m*.

##### Two temporal channels model

For the grid fit, we generated a 10 by 10 grid for the sustained and the transient weight (from 10^-5^ to 1 with 10 equal steps). Then we implemented the search fit step as above, with lower bound [0, 0] and upper bound [1, 1]. See *trf_gridFit.m* and *trf_fineFit.m*.

##### Linear model

The linear model does not require a search or seeds. Instead, we fit the 100 bootstrapped data sets per ROI by linear regression, giving us 100 estimates of the gain factor, *g*, per ROI.

#### Statistical analysis

We compared model accuracy of the CTS and the linear model. Because the models have different numbers of free parameters, it is important to obtain an unbiased estimated of model accuracy, which we did by leave-one-out cross validation. For each ROI, and for each of the 100 bootstrapped sets of Beta-weights, we fit 13 linear models and 13 CTS models by leaving out each of the 13 temporal stimuli. For each bootstrap, we thus obtained 13 left-out predictions, which were concatenated and compared to the 13 Beta-weights by coefficient of determination, R^2^:

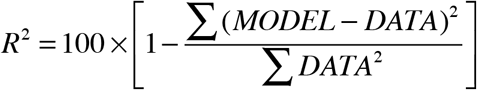

This yielded 100 R^2^’s per ROI, and we summarized model accuracy as the median and 50% confidence interval derived from these 100 values.

Note that the coefficient of determination, R^2^, is bounded by [-∞, 1], as the residuals between model and data can be larger than the data. In contrast, r^2^ is bounded by [0, 1].

##### Noise ceilings

The noise ceiling represents the highest accuracy a model can achieve given the signal-to-noise ratio in the data, irrespective of the specific model used. We computed noise ceilings stochastically based on the mean and standard error of the GLM-Beta weights from bootstrapped estimates, as implemented in (Kay et al., 2013a).

##### Flat model

We computed a model that assumes the BOLD responses to all stimuli are identical (‘flat model’) as a further basis of comparison to the CTS and linear models. Like the CTS and the linear model, the accuracy of the flat model was computed by leave-one-out cross validation. (The cross-validated predictions from the flat model are not quite identical across conditions, because the mean is affected by the left-out data.)

##### Parameter Recovery

To estimate how well model parameters are specified, for each visual area, we simulated the CTS model responses by first generating the predicted fMRI Beta-weight for each temporal condition. The parameters used for simulation were the median of each of the parameters from the bootstrapped fits to the data. We then added noise to each Beta-weight by randomly sampling from a normal distribution whose standard deviation was matched to the standard error in the bootstrapped data, averaged across the temporal conditions. We added noise 1,000 times per ROI, and then solved the CTS model for the 1,000 simulated responses using the same procedure used with the actual data. The parameters recovered from this fitting procedure provide an estimate of how well specified each parameter is given the form of the model (including the parameters) and the noise level in the data.

### 2.3 Public Data Sets and Software Code

To ensure that our computational methods are reproducible, all data and all software are made publicly available via an open science framework site, https://osf.io/v843t/. The software repository includes scripts of the form *trf_mkFigure2* to reproduce figure 2, etc., as in prior publications (e.g., Winawer and Parvizi, 2016).

## 3 Results

### 3.1 Measuring temporal summation in visual cortex

In each trial of the experiment, participants viewed either one or two pulses of a static spatial contrast pattern. The pattern was an independently generated band-pass noise image (24° diameter), used in prior studies of spatial encoding (Kay et al., 2013a; Kay et al., 2013b). For the two-pulse trials, the two spatial patterns were identical. Each trial used one of thirteen distinct time courses (Figure 2A). The durations of the one-pulse stimuli and the ISIs of the two-pulse stimuli were the same: 0, 1, 2, 4, 8,16, 32, or 64 video frames of a 60 Hz monitor (i.e., 0, 17, 33, 67, 134, 267, 533 ms). Each pulse in the 2-pulse stimuli was 8 frames (134 ms). The 0-ms one-pulse stimulus was a blank (mean luminance), and the two-pulse stimulus with 0 ISI was identical to the one-pulse stimulus of twice the length (16 frames, 267 ms). Four participants were scanned, and data were binned into nine bilateral, eccentricity-restricted (2-10°) visual areas defined from a separate retinotopy scan (Figure 2C).

The fMRI data were analyzed in two stages. First, we extracted the amplitude (Beta-weight) for each of the 13 temporal conditions using a variation of the general linear model, “GLM denoise” (Kay et al., 2013c), a technique that improves the signal-to-noise ratio by including noise regressors in the GLM (Figure 2B). Second, we fitted the temporal pRF model to the GLM Beta-weights, averaged across voxels within ROIs.

### 3.2 Temporal summation in visual cortex is subadditive

We tested the linearity of the fMRI BOLD signal in each visual area. To do so, we assume a time-invariant linear system such that the BOLD amplitude (GLM Beta-weight) is proportional to the total stimulus duration within the trial. Due to the linearity assumption, the form of the neural impulse response function does not affect the pattern of the predicted BOLD amplitudes. For example, the linear prediction is that a stimulus of duration *2t* produces twice the amplitude as a stimulus of duration *t,* and the same amplitude as a two-pulse stimulus, with total duration 2*t* (Figure 3A). This prediction is not borne out by the data. The response to a stimulus of length *2t* is about 75% of the linear prediction in V1 and 50% in area TO (a homolog of macaque areas MT and MST) (Figure 3B, left panel). This systematic failure of linearity is found in all visual areas measured, with temporal summation ratios below 0.8 for all ROIs, and a tendency toward lower ratios in later areas (Figure 3C). The BOLD amplitudes to all stimuli are low (<1%) because the stimuli are brief, compared to measurements of visual cortex using moving stimuli or a block design with multiple static stimuli, where percent BOLD changes can be several percent.

**Figure 3.**
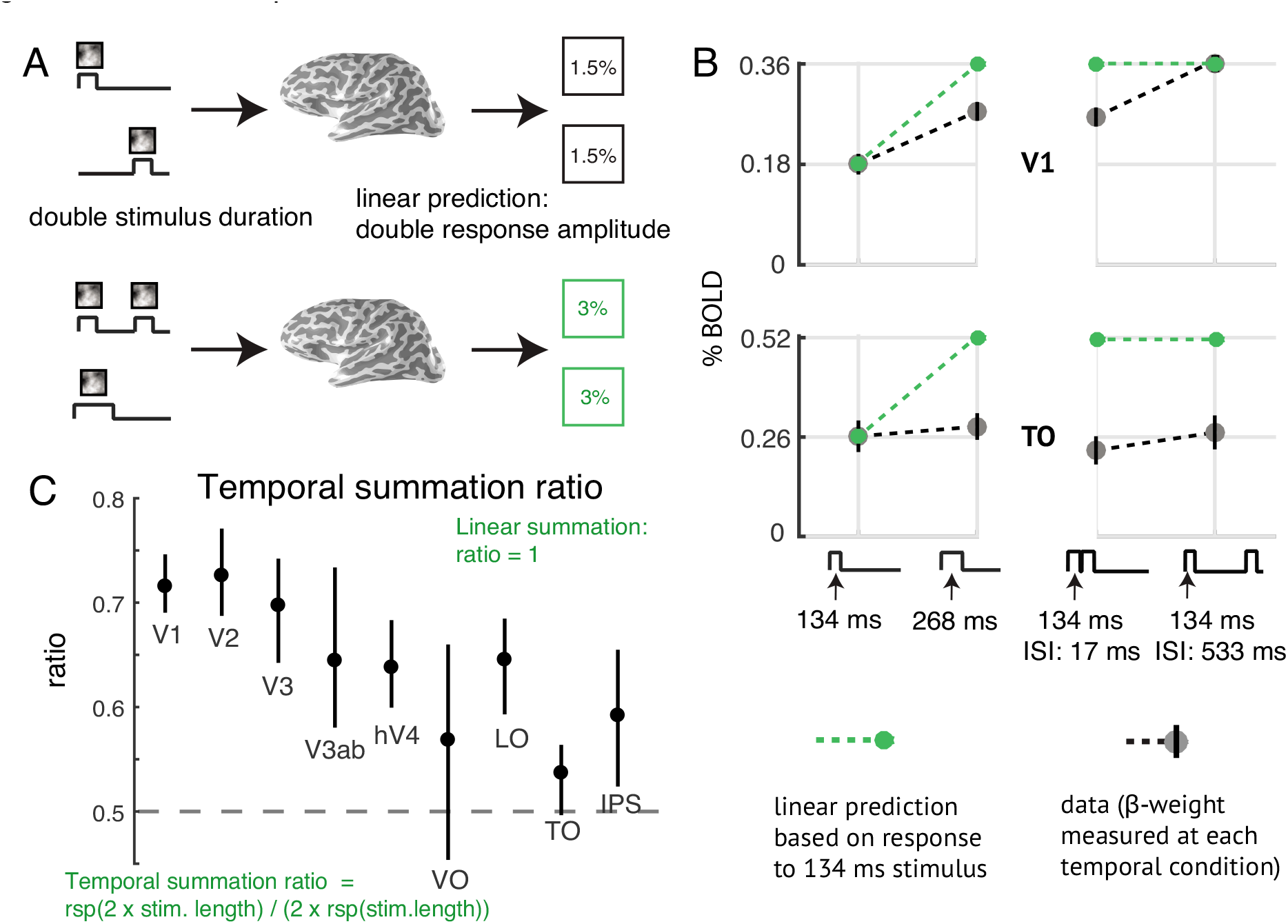
Sub-linear temporal summation in visual cortex. *(A) Linear temporal summation prediction*. The sum of the responses to two separate events (top) is equal to the response to the two events in the same trial, with or without a brief gap between them (bottom). *(B) Sub-linear temporal summation*. Gray circles are the measured responses to a 134-ms pulse, a 268-ms pulse, and two 134-ms pulses, with either a 17-ms or 134-ms gap between pulses. Plots show the mean across participants and 50% CI (bootstrapped across repeated scans within each participant). The green circles and dotted lines are the linear prediction based on the response to the single 134-ms pulse. For V1, the measured responses are less than the linear prediction except when there is a long gap. For area TO, all responses are less than the linear prediction*. (C) Temporal summation ratio*. Temporal summation ratio is the response to a stimulus of length 2x divided by twice the response to a single pulse stimulus of length x, averaged across 5 stimulus pairs (17 and 34 ms, 34 and 68 ms, etc.). Linear summation occurs when the temporal summation ratio is 1. The temporal summation ratio is less than 1 in all visual areas, indicating compressive temporal summation. The ratio is higher in early visual areas (~0.7 in V1-V3), and lower in later areas (~0.5 to 0.65). Error bars represent the 50% CI (bootstrapped across scans). The ROIs on the x-axis are arranged in order of increasing spatial pRF size at 5 deg eccentricity, as a proxy for order in the visual hierarchy. Figure made from the script trf_mkFigure3.m.

A further failure of linearity occurs for trials with two pulses and variable ISI: the response is larger when the ISI is longer, especially in V1. The linear prediction is that the amplitudes are the same, and double the response to the one-pulse stimulus (Figure 3B, right). When the ISI is relatively long (528 ms), the response in V1 is close to the linear prediction made from the one-pulse stimulus. In TO, even with a long ISI the response is still well below the linear prediction. This pattern, whereby the response to a second stimulus is reduced for short ISIs, and larger for longer ISIs, is often called adaptation and recovery (Priebe et al., 2002; Kohn, 2007). For TO, the recovery time is longer than V1, and longer than the longest ISI we tested.

### 3.3 The temporal subadditivity is captured by a compressive temporal summation model (CTS)

We modeled the temporal subadditivity with a compressive temporal summation model (“CTS”) (Figure 4A). The CTS model has a Linear-Nonlinear (LN) structure. The linear stage convolves the stimulus time course with a temporal impulse response function (parameterized by the time constant τ). The nonlinear stage passes the linear output through a static nonlinearity, divisive normalization. The normalization is implemented by squaring the linear response at each time point (as in (Heeger, 1992)), and dividing this by the sum of two terms, a semi-saturation constant (σ) and the linear response, both of which are squared (Heeger, 1992). Squaring is widely used for modeling neural computations such as color (Helmholtz, 1886; Koenderink et al., 1972) and motion (Adelson and Bergen, 1985; Simoncelli and Heeger, 1998). The normalization model was developed to describe responses at the level of single neurons. However, we can generalize it to an fMRI voxel by assuming that the neurons within a voxel share a normalization pool, and the voxel sums across neurons within it. In this case, the normalization equation has the same term in the numerator and denominator, as implemented in the CTS model. (See also: Relationship to Divisive Normalization in (Kay et al., 2013a).)

**Figure 4.**
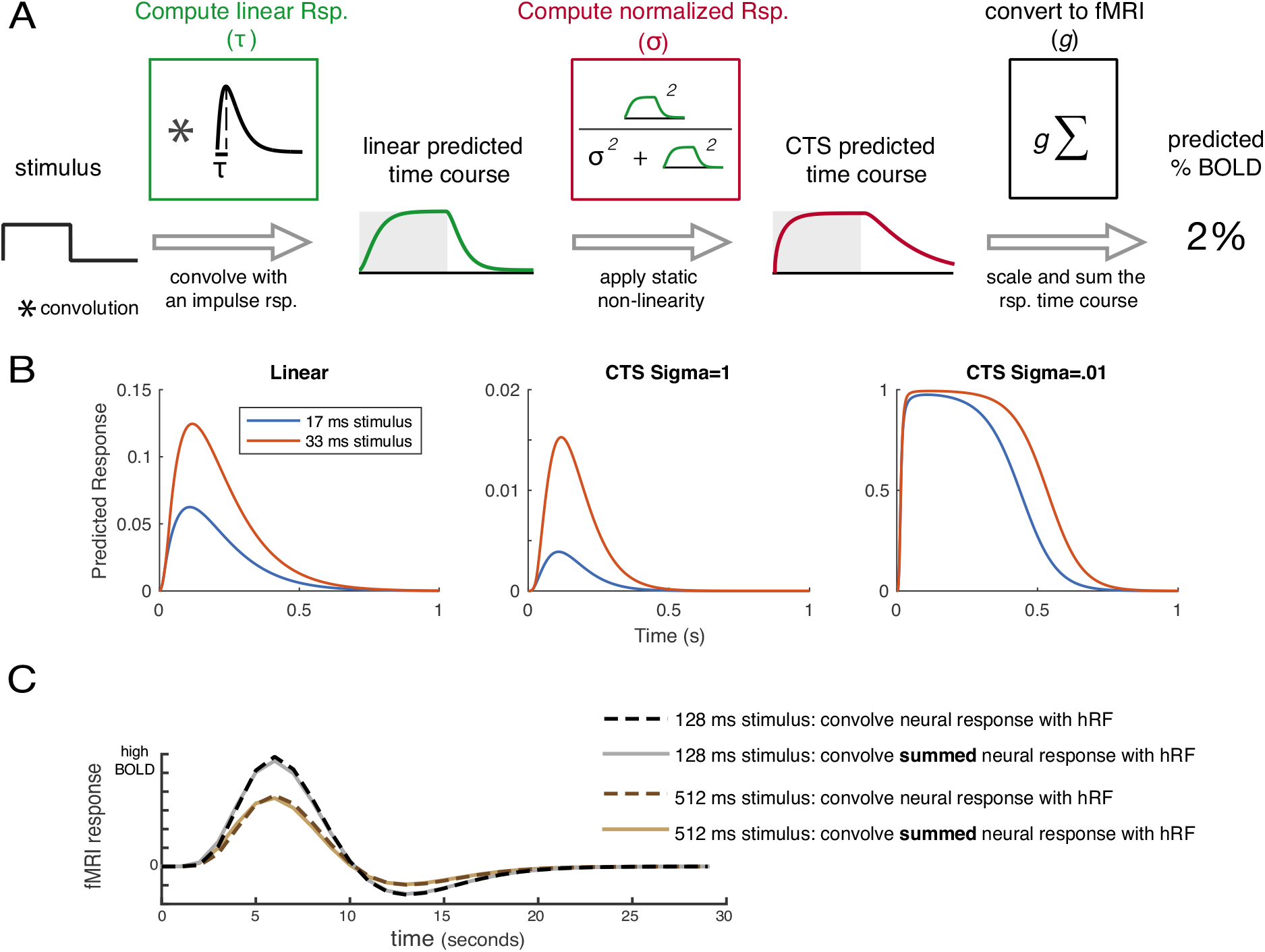
Compressive temporal summation (CTS) model. (A) The CTS model. The CTS model takes the binary stimulus time course for a trial as input (1 whenever the contrast pattern is present, 0 whenever it is absent). The input is convolved with an impulse response function parameterized by τϵto produce a linear prediction. The second stage is a divisive normalization computation. The numerator is the linear prediction raised point-wise and squared. The denominator is the sum of the squared semisaturation constant, σ^2^, and the squared linear response. Finally, the time-varying CTS prediction is summed and scaled (g) to predict the percent BOLD response. The CTS model was fit for each ROI by finding the values of τ, σ, and g that minimized the squared error between the model predictions and the GLM Beta-weights. (B) Predicted neural time series. The left panel shows predictions from a linear model to a 17-ms and 33-ms stimulus, assuming an impulse response function with time constant 100ms. The other two panels show the CTS model predictions with large σϵ(middle) and small σϵ(right). When σϵis small, the predicted time series are similar for the two stimuli. (C) CTS-predicted BOLD time series. CTS model predictions were computed for two stimuli, a 128-ms and a 512-ms single pulse stimulus (assuming τϵ= 0.05; σϵ= 0.01). The CTS model predictions were then passed through a hemodynamic response function (HRF) in one of two ways, either by convolving the CTS model prediction with the HRF (dashed lines), or by convolving the HRF with a single number for each stimulus, the sum of the CTS model prediction (solid lines). For each stimulus, the predicted response to the summed CTS response and to the time-varying CTS response is almost identical. Further, the BOLD response to the longer stimulus is the same shape as the response to the briefer stimulus, just scaled in amplitude.

We illustrate the effect of the CTS model with example responses to brief stimuli (17 ms and 33 ms), assuming an impulse response function with time constant 100-ms (Figure 4B). For a linear model, the predicted response to the longer stimulus peaks at almost double the value of the briefer stimulus. For the CTS model with a large σ, the response more than doubles for the long stimulus compared to the brief stimulus, due to the squaring in the numerator. When σϵis small, the model is compressive, as the response to the longer stimulus is very similar to the brief stimulus.

To relate the CTS model output to the BOLD signal, we summed the predicted CTS output for a trial, and scaled this by a gain parameter, *g*, to convert to units of percent BOLD change. We sum the CTS output to give a single value per temporal condition, which can be compared to the Beta-weight in each condition, fit from the GLM. If we instead convolve the time-varying CTS model prediction with an HRF, rather than convolving the summed CTS model prediction with the HRF, the predicted BOLD response is nearly identical (Figure 4C). We note that while we refer to the model as compressive, technically the normalization model amplifies the output when the instantaneous linear response is low due to the squaring in the numerator, and compresses the response when the amplitude is high. However, for all temporal conditions we tested, the model output is compressive in the sense that the predicted response for any of our single pulse stimuli is less than the linear prediction from a briefer stimulus.

We compared the CTS model to a linear model by measuring cross-validated accuracy. The CTS model is more accurate than the linear model for all areas (Figure 5). The linear model substantially underpredicts responses to short durations and overpredicts responses to long durations, whereas the CTS model does not. Further, the predictions of the linear model do not depend on ISI, whereas the CTS model correctly predicts that the response amplitude increases with longer ISI. The cross-validated CTS model predicts the left-out fMRI responses with accuracy between 81% and 98% across the 9 ROIs. This represents a large improvement compared to the linear model for every area (Figure 5B). The improvement is especially pronounced in later than early areas (LO/TO/IPS vs. V1-V3).

**Figure 5.**
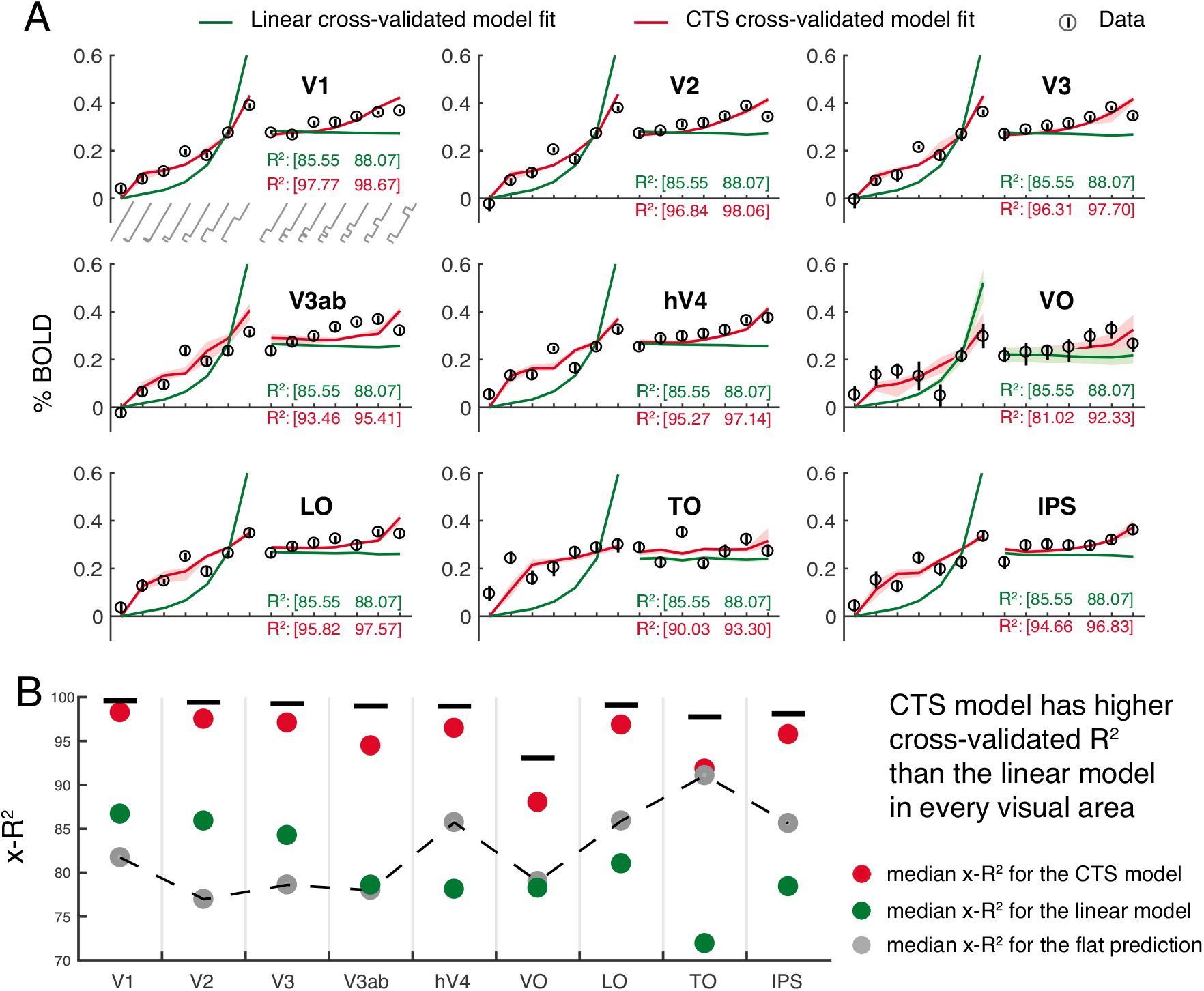
CTS model fits to BOLD data across visual areas. *(A) Data and predictions*. BOLD responses to each temporal condition averaged across participants are plotted as circles. The temporal conditions on the x-axis show increasing durations of one-pulse stimuli (0 to 533 ms; left) and increasing ISI of two-pulse stimuli (0 to 533 ms, right). Stimulus temporal conditions are as in Figure 2A. Error bars show the 50% CI bootstrapped across repeated scans. Predictions for the linear (green) and CTS (red) model fits are computed by leave-out-one-condition cross-validation. Shaded regions represent the 50% CI of the predictions across bootstraps (not visible for most of the linear fit because the CI is narrow). *(B)* The cross-validated accuracy (x-R^2^) is higher for the CTS model, compared to the linear model in each area. Each circle represents the median cross-validated R^2^ for each model and the error bar is the 50% CI across bootstraps. (Figure made from the script trf_mkFigure5.m.)

The CTS model is also more accurate than a flat model (Figure 5B). The flat model predicts the same response amplitude to all stimuli. This indicates that although the BOLD responses are relatively small (low percent signal change) and compressive (similar for different duration stimuli), there are nonetheless meaningful differences in the response amplitudes to different temporal conditions. Importantly, the CTS model accurately captures these differences, as it is substantially more accurate than the flat model. One notable exception is area TO, where the BOLD responses are most compressive: here the CTS model is only slightly more accurate than the flat model (and both are much more accurate than the linear model). In contrast, the linear model is more accurate than the flat model only in early visual areas (V1-V3) and less accurate in higher visual areas.

We note that although the cross-validated accuracy of the CTS model is high (close to the noise ceiling in all areas), some data points appear to deviate systematically from the model predictions – for example, the response to the 17-ms single-pulse stimulus is under-predicted in TO, and the 67-ms single pulse stimuli are under-predicted in multiple areas. We do not try to interpret these particular data points as they were not robust to replication (section 3.6).

### 3.4 The CTS model fits capture systematic differences between areas

The CTS model is parameterized by τ, σ, and a gain factor, *g*. τϵis the time to peak in the temporal impulse response function, and therefore is related to temporal summation window length; σϵis the semisaturation constant, and reflects how much the CTS prediction deviates from the linear prediction. When σϵis lower, the response is more compressive. The intuition for this is that when σϵis small, the numerator and denominator have similar values, so that the model output is relatively invariant to stimulus duration (hence more compressive). In contrast, when σϵis very large, the denominator is approximately a constant, so there is little normalization. In later visual areas (hV4 - IPS), σ is about 10 times lower than earlier areas (V1 – V3ab) (~0.003 v ~0.03; Figure 6A, right), consistent with temporal summation being more sub-linear in the model-free summary (Figure 3C). The more pronounced sublinearity later in the visual hierarchy is qualitatively similar to the pattern found for spatial summation (Kay et al., 2013a). A consequence of more compressive temporal summation is that the response amplitude varies less with minor changes in stimulus duration, just as greater compression of spatial summation predicts more tolerance to changes in size and position (Kay et al., 2013a). From the current fMRI data set, there is also a tendency toward shorter time constants (τ) in earlier areas, with some exceptions (except for VO, V1-V3 have the smallest τ, ~50 ms; Figure 6A).

**Figure 6.**
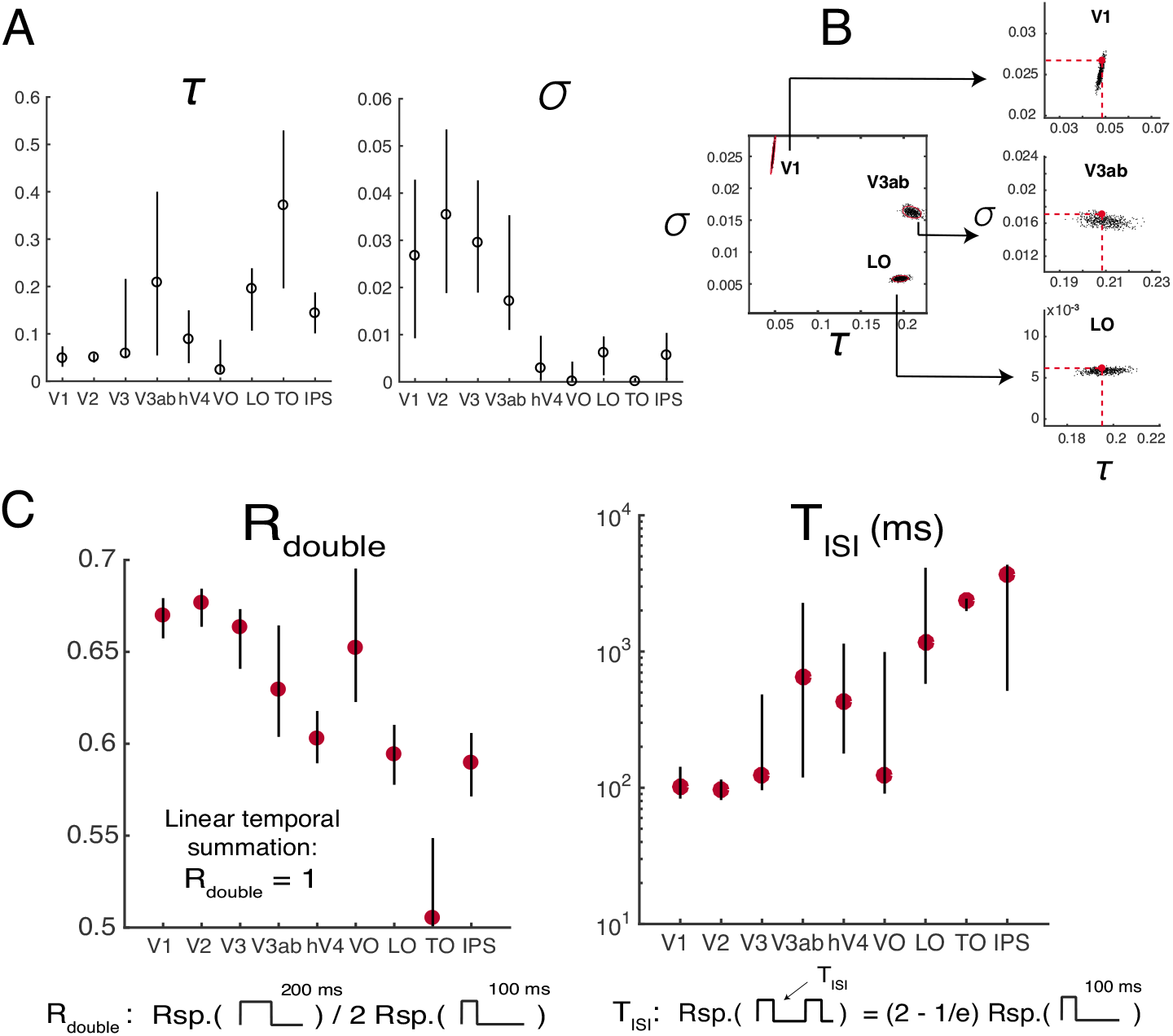
CTS model parameters and summary metrics. *(A) CTS model parameter estimates*. The estimated parameter σϵis smaller for later (~0.003, hV4 - IPS), compared to earlier visual areas (~0.3, V1 – V3ab), indicating that the temporal sum in the later visual areas deviates more from the linear sum. The time constant τϵis short in V1-V3, compared to most of the later visual areas. *(B) CTS parameter recovery*. The precision with which parameters can be fit depends on the noise level in the data and the specific parameter values. We simulated data using the median τϵand σϵfrom V1, V3ab, and LO, and the noise levels estimated from these areas. The analysis shows that under these conditions, τ (x-axes) is specified most precisely in V1 and least precisely in V3ab; the opposite pattern is found for σ (y-axes). *(C) Summary metrics*. Two summary metrics of the CTS model reveal a pattern across ROIs. Rdouble is the ratio of the predicted response to a 200-ms pulse divided by twice the response to a 100-ms pulse. Rdouble is below 1 for all ROIs, indicating sub-additivity, and decreases along the visual hierarchy (V1-V3, ~0.67, LO-IPS, < 0.6). TISI is the length of ISI required for the response to two 100-ms pulses to approach the linear prediction. TISI is short in the earlier areas (V1-V3, ~100 ms) compared to most of the later areas. (Figure made from the script trf_mkFigure6.m.)

The precision of our parameter estimates in each area depends on the BOLD noise level (the confidence interval of the Beta-weights), as well as the specific parameters estimated for that area. To understand how these factors interact, we simulated 1,000 data sets for each of 3 areas – V1, V3ab, and LO. The simulations used the median parameter fits for each area (τϵand σ) to generate a noiseless prediction. We then added noise independently for each of the 1,000 predictions, according to the noise level in the fMRI measures for that area. Finally, we solved the CTS model for each of the predicted set of responses and analyzed the parameters. This parameter recovery analysis reveals two important results. First, it shows that the parameters for the different areas are distinguishable: models solved from simulations matched to V1, say, are not confusable with models solved from simulations matched to V3ab or LO (Figure 6B). Second, the analysis shows that the precision of the parameter estimates differs across areas. For example, for V1, τϵis more precisely specified than σ, whereas for LO, σϵis more precisely specified than τϵ(Figure 6B, insets). V3ab is intermediate. These simulations are consistent with the observation that model solutions on the bootstrapped data show a smaller confidence interval for τϵthan for σϵin V1, and the reverse for LO (Figure 6A).

To further examine the differences in temporal processing between ROIs, we summarized the CTS model predictions to each ROI response in terms of two metrics that have more directly interpretable units: R_double_ and T_ISI_ (Figure 6C). R_double_ is the ratio between the CTS-predicted BOLD response to a 100-ms stimulus and a 200-ms stimulus. Lower R_double_ means more compressive temporal summation. Later visual areas have lower R_double_ than earlier ones. T_ISI_ is the minimal duration separating two 100-ms pulses such that the response to the paired stimuli is close to the linear prediction from the single stimulus. Similar to previous measurements at longer time scales (Weiner et al., 2010; Mattar et al., 2016), the recovery time is longer for later than earlier visual areas.

### 3.5 The CTS parameters do no vary systematically with eccentricities from 2 to 12 degrees

Prior work has shown that temporal encoding in V1 differs between fovea and periphery (Horiguchi et al., 2009). In a separate analysis, we asked whether the CTS model parameters differed as a function of eccentricity. We did not find reliable differences for parafovea (2º-5º) versus periphery (5º-10º), either in the response amplitude (Figure 7A) or in the summary metrics (Figure 7B). This may be due to the limited range of eccentricities: Horiguchi et al (2009) found the biggest difference in temporal sensitivity between fovea and the far periphery (20º-60º), with only minimal differences between the low-to-mid eccentricity bins we tested.

**Figure 7.**
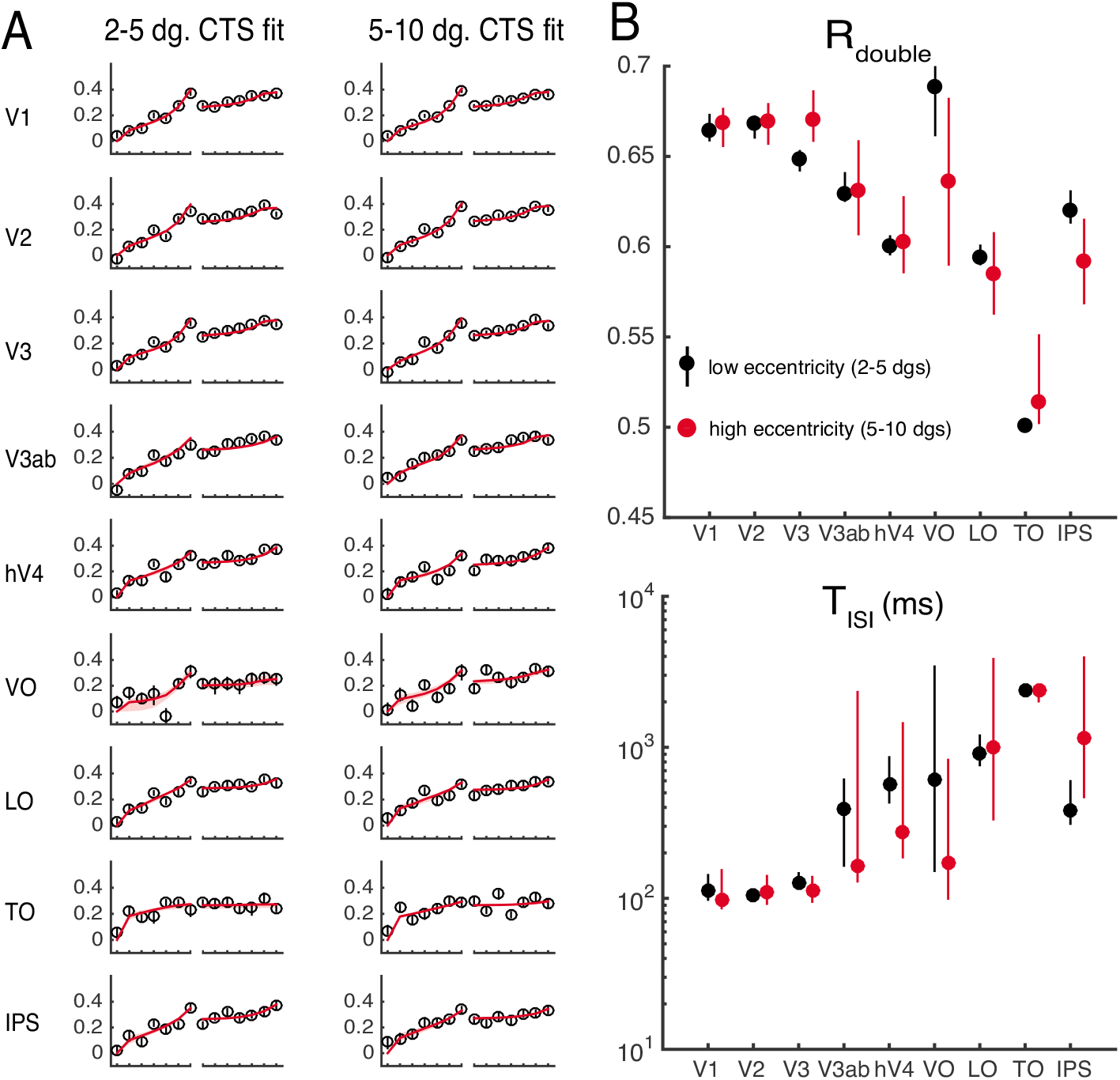
CTS model fits by eccentricity. Data from the main fMRI experiment are re-plotted separating each ROI into 2 eccentricity bins. *(A) CTS model fit to low and high eccentricity bins*. The left panels are the data and CTS model fits restricted to voxels with population receptive field centers within 2 - 5°. The right panels are data and CTS model fits restricted to voxels within 5 - 10° eccentricity. *(B) Summarized metrics for different eccentricity bins*. The summarized metrics do not differ systematically between the two eccentricity ranges. Each dot represents the median of the metrics summarized for 100 bootstraps of data (across scans), error bars represent 50% confidence interval. (Figure made from the script trf_mkFigure7.m.)

### 3.6 The CTS model fits replicate across experiments

We conducted a separate experiment with the identical temporal profiles and two different classes of images – pink noise and faces embedded in pink noise (Figure 8A). This experiment tests the generalizability across spatial pattern. Because faces were used as one of the textures in this experiment, we included an additional region of interest – a face-selective area, which is a combination of the occipital face area (OFA) and the fusiform face area (FFA). A single model was fit to each ROI for each participant, assuming independent gain parameters for the two stimulus classes, and the same time constant and semi-saturation constant. The results from the main experiment hold: all visual areas in the second experiment sum sub-linearly in time, with the CTS model fitting the data more accurately than the linear model (Figure 8B). Moreover, as with the main experiment, later areas tended to sum more sub-linearly compared to the earlier ones (Figure 8C). The response amplitudes are slightly lower than those in the main experiment due to stimulus selectivity, and the responses are noisier due to fewer trials per condition; otherwise the pattern of responses is highly similar.

**Figure 8.**
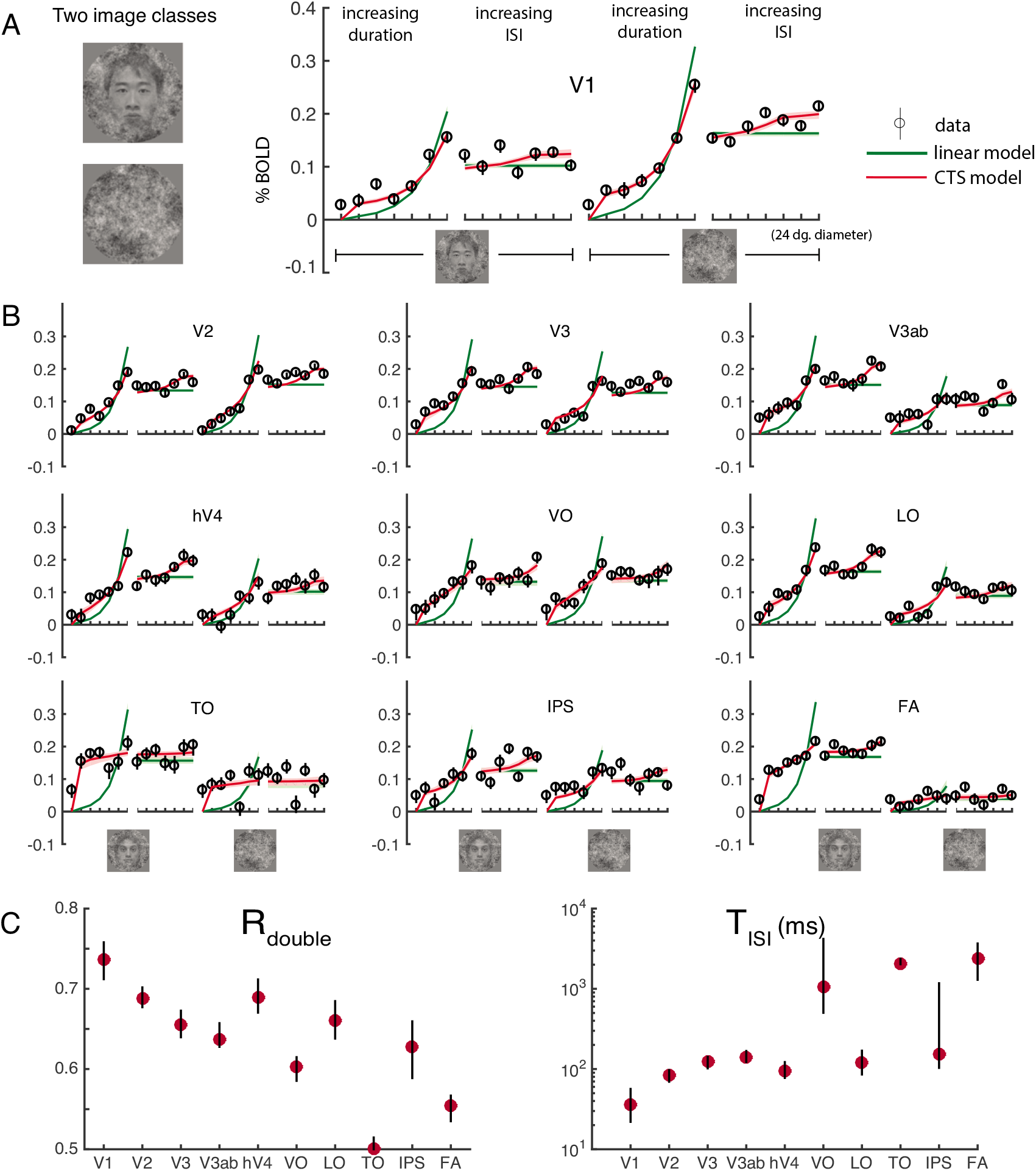
FMRI data and model fits from a second experiment. (A) Stimuli and V1 responses. The two stimulus classes – noise patterns and faces embedded in noise patterns, were randomly interleaved within runs. Temporal conditions were identical to those in Figure 4. The general pattern of responses and model fits are highly similar to those in the main experiment, with the CTS model fitting the data much more accurately than the linear model. (B) CTS model fit to extrastriate visual areas. The CTS model (red) fit the data more accurately than the linear model in all visual areas. (C) Parameters derived using the CTS model fit. The derived metrics, R_double_ and T_ISI_ show similar patterns as in the main experiment: decreased R_double_ and increased T_ISI_ in higher visual areas. (Figure made from the script trf_mkFigure8.m.)

### 3.7 Differences in parameters across ROIs are not explained by differences in HRFs

We found that the CTS parameters representing temporal processing differed systematically along the visual hierarchy, with a tendency towards a pronounced nonlinearity and longer time constant in later visual areas. These results were obtained with a model in which a single HRF was fitted to each individual subject, but not to each ROI separately. Fitting a single HRF to each area is robust in that it reduces sensitivity to noise within an area as signal). However, if the actual HRF differs systematically across areas, then it is possible that enforcing a single HRF will result in biased estimates of the Beta-weights. In this section, we consider the possibility that the differences in derived metrics (R_double_ and T_ISI_) across ROIs might be explained by variations in the HRF rather than differences in the underlying neuronal responses.

**Figure 9.**
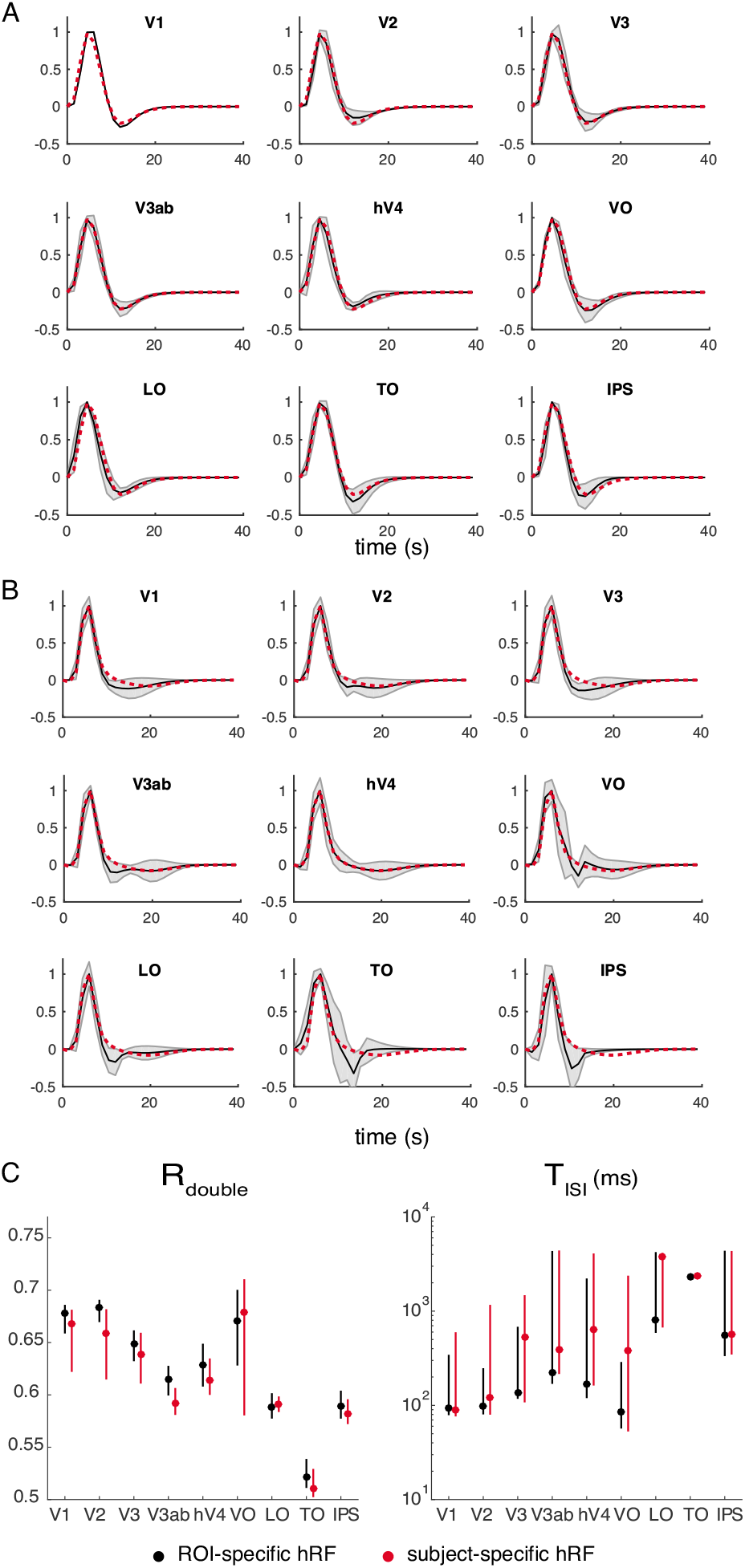
HRFs for individual ROIs. *(A)* HRFs derived from the retinotopy experiment. The shaded black curves represent the estimated HRF from each ROI (mean and std. across subjects). The dotted red line represents the mean HRF estimated for hV4, replotted in each panel for comparison. The hV4 HRF does not differ much from the other individually estimated HRFs. *(B)* Same as panel A, but the HRFs are derived from the temporal experiment. *(C)* CTS summary metrics (R_double_ or T_ISI_) derived from the CTS model, fit either with subject- and ROI-specific HRFs (black) or only subject-specific ROIs (red). (Figures made from the script trf_mkFigure9.m.)

To address this question, we estimated a separate HRF for each ROI and each subject. We estimated the ROI-specific HRF in two ways: from the retinotopy experiment and from the first temporal experiment. For each area, the HRFs were parameterized as a difference of two gamma functions (Friston et al., 1998; Worsley et al., 2002). The resulting HRFs were broadly similar across ROIs. For example, the time course of the HRF of an intermediate area, hV4, is within a standard deviation of the time course of all other ROIs at almost all time points from both retinotopy and the temporal experiment (Figure 9A,B). There are some qualitative differences, such as a larger post-stimulus undershoot in later areas, particularly as estimated by the temporal experiment (VO, LO, TO, IPS). To assess the impact of these modest differences in the HRFs across areas, we recomputed the CTS model parameters and the two derived summary metrics, R_double_ and T_ISI_ (Figure 9C). The general pattern of results is the same whether the HRFs are fit to each area or to each individual: R_double_ tends to decrease along the visual hierarchy, and TISI increases.

### 3.8 The CTS model is more accurate than a two temporal channels model in later visual areas

The CTS model was implemented to capture subadditive temporal summation using canonical neural computations (filtering, exponentiation, normalization). An alternative model, in which neuronal responses are thought to reflect the outputs of two temporal channels, has been proposed to account for psychophysical temporal sensitivity (Watson and Robson, 1981; Hess and Plant, 1985; Watson, 1986) and fMRI responses in V1 (Horiguchi et al., 2009) and extrastriate cortex (Stigliani et al., 2017). This two-temporal-channels model linearly combines the output from a sustained and a transient temporal frequency channel. The sustained channel has a mostly positive IRF and is linear and the transient channel has a balanced (zero-sum) IRF and its output is squared (Figure 10A). The two temporal channels model contains filtering and exponentiation but not normalization. The specific forms of the impulse response functions in the two temporal channels model are derived from psychophysics, not neural data, and hence are assumed to be the same in all visual areas; the model is fit only by varying the relative weights of the two channels.

**Figure 10.**
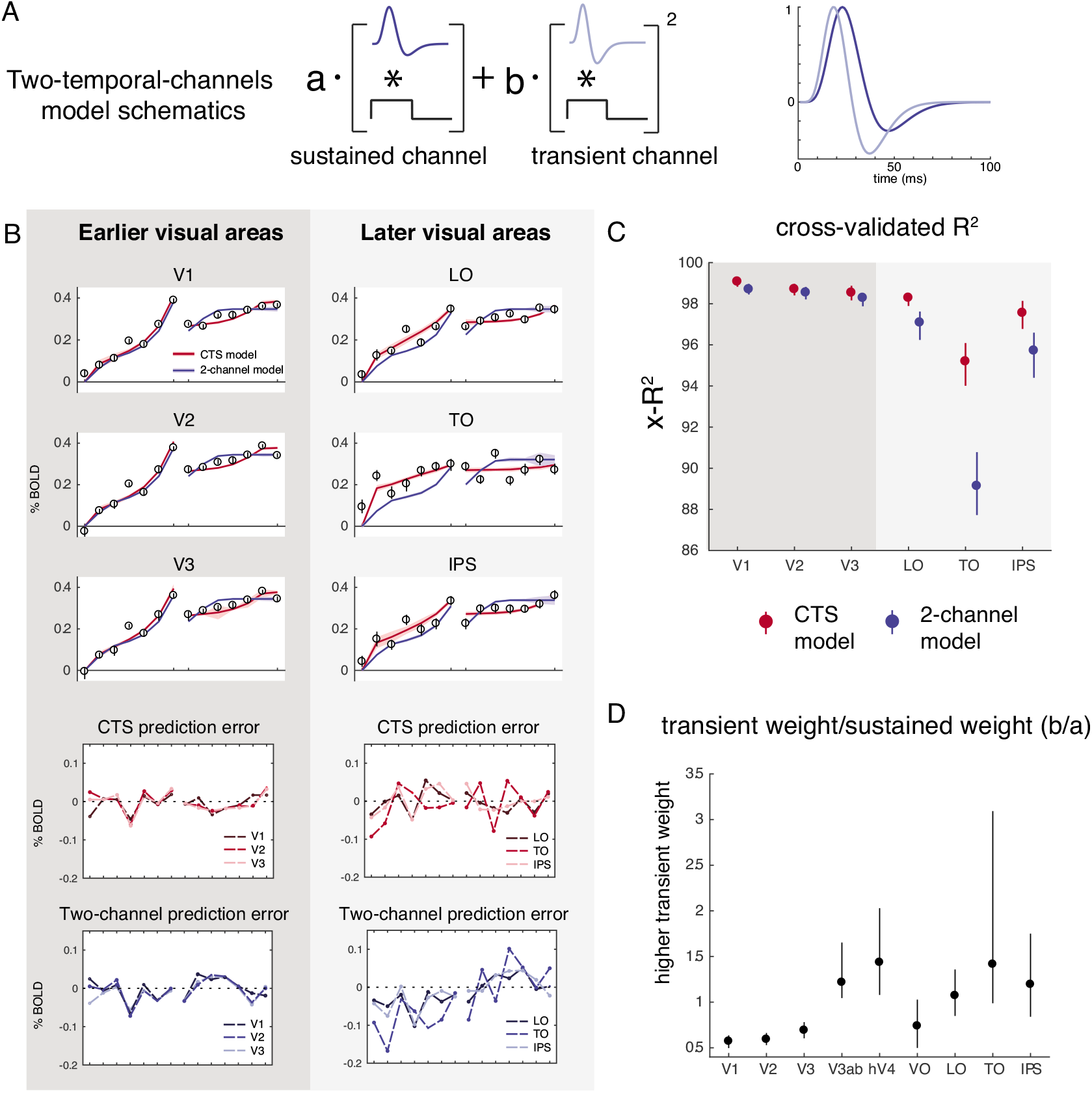
The CTS model describes data in later visual areas better than the two temporal channels model. *(A)* Two temporal channels model schematic. The model combines the outputs from a sustained channel (mostly positive IRF, linear output) and a transient channel (biphasic IRF, squared output). This model was used by Horiguchi et al. (2009) to fit FMRI data to temporal modulations in luminance. The fMRI implementation was adapted from previous models that account for psychophysical data (Watson, 1986). *(B)* The CTS and two-temporal-channels model fit the data about equally well in V1-V3, whereas the CTS model fits the data better in later areas (LO, TO, and IPS). In the later visual areas, the two-temporal-channels model systematically under-predicts the response to the one-pulse stimulus and over-predicts the response to two-pulse stimuli (lower four panels). *(C)* The cross-validated accuracy (x-R^2^) is higher for the CTS model in later areas. (D) The predicted weights for the sustained versus the transient channels differ across areas. The ratio between the weights for the transient versus for the sustained channels is plotted for each visual area (median, and 50% CI across bootstraps). Later visual areas tend to show increasingly high transient weights, consistent with Stigliani et al., 2017. (Figure made from the script trf_mkFigure10.m.)

We fit the two-temporal-channels model to the bootstrapped Beta-weights estimated from the first temporal experiment, and compared this to the CTS model fits. In early visual areas (V1 – V3), the two models have similar accuracy (as assessed by cross-validated R^2^). In later visual areas (for example, in LO, TO and IPS), the CTS model captures the data better. For the later areas, the two-temporal-channels model systematically under-predicts Beta-weights for the one-pulse conditions, and overpredicts the two-pulse conditions (Figure 10B–C): Because of the relatively brief time scales of the IRFs in the two-temporal-channels model, the predicted response to the second of two pulses is largely unaffected by the first pulse. This will result in an over-prediction for any visual area with long temporal windows. As noted earlier, measurements further in the periphery of V1 have greater sensitivity to stimulus transients (Horiguchi et al., 2009); it is therefore likely that had our measurements extended into the far periphery, the CTS model would need to be augmented with a second, transient channel.

### 3.9 Alternative implementations of CTS model

The nonlinear component of the CTS model was implemented as a divisive normalization. This model fit the data much more accurately than a linear model, and divisive normalization is a good descriptor of a wide range of neural phenomena (Carandini and Heeger, 2012). However, there are many choices of static nonlinearities. In prior work, a power law static nonlinearity was used to model compressive spatial summation in fMRI (Kay et al., 2013a; Winawer et al., 2013). Although the form of the two nonlinearities differ, we found that refitting the fMRI data with the power law nonlinearity produced results that were highly similar to the fits with the divisive normalization implementation (Figure 11). The model accuracy was not distinguishable when using a power law vs divisive normalization, and the two derived metrices, R_double_ and T_isi_, showed the same pattern. Hence the fMRI data in these experiments do not distinguish the two forms of the compressive nonlinearity. The power law nonlinearity has the advantage of ease of interpretation – the exponent indicates how much the response deviates from linear. Divisive normalization has the advantage of strong support from many neural systems (Carandini and Heeger, 2012). It might be possible to distinguish the two by measuring responses to stimuli with very brief durations or very low contrasts. One difference between the two nonlinearities is the precision in which parameters are recovered. For example, τ is recovered with high precision for most visual areas in the divisive normalization implementation; for the power-law implementation, the exponent εϵis recovered more accurately than τϵ(simulations not shown).

**Figure 11.**
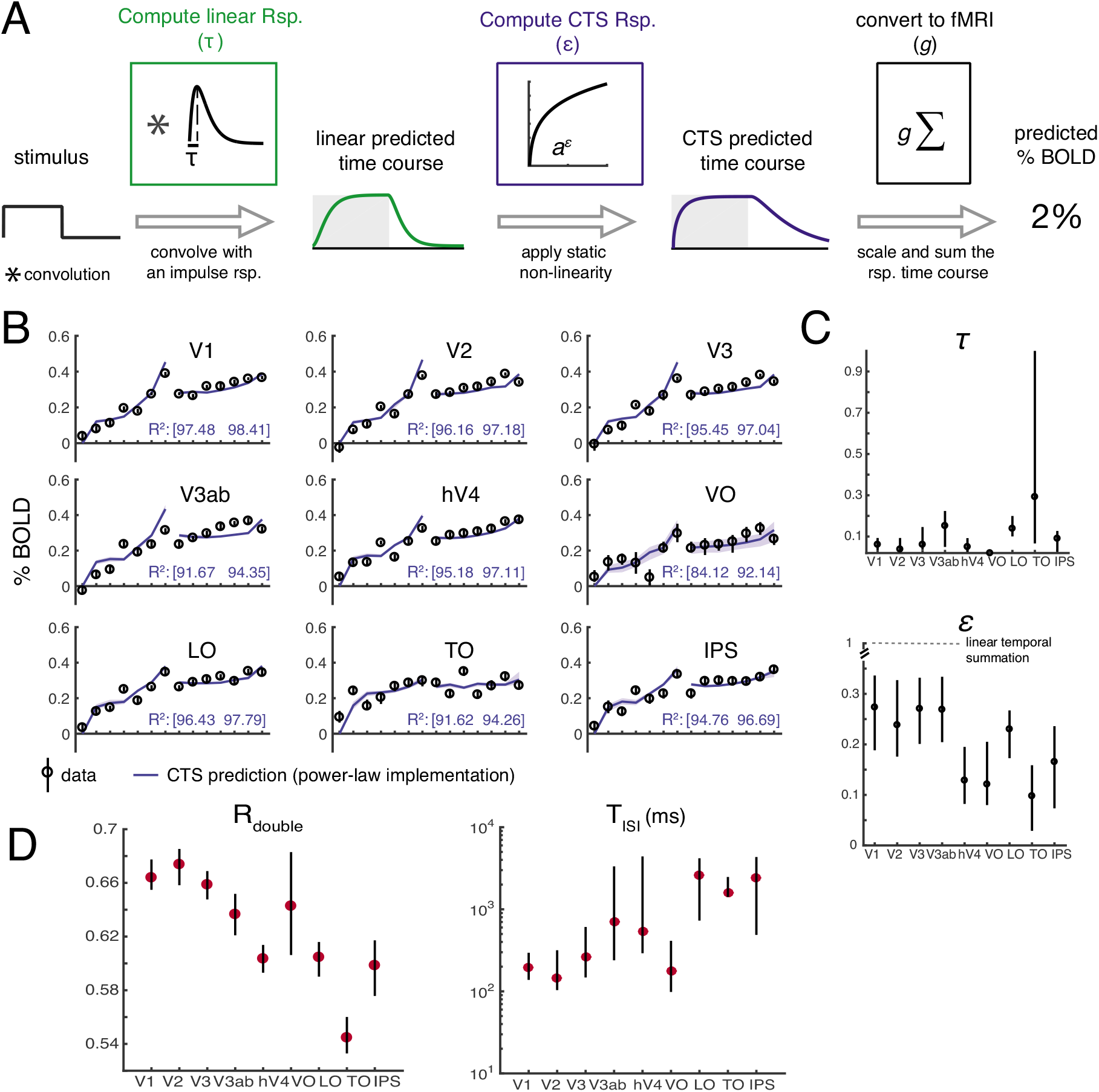
CTS model implemented with a power-law nonlinearity. *(A) Model description*. The model is identical to that shown in Figure 4, except that the static nonlinearity is a power-law (parameterized by ε) rather than a divisive normalization. If εϵis 1, the model is linear, and if εϵis less than 1, it is compressive. *(B) Cross-validated model fit to the data from the main fMRI experiment*. The model describes the data with high accuracy. See figure 5A for plotting conventions. *(C) Model parameters*. The estimated exponent εϵis below 1 in each area, and is lower (more sub-linear) in later areas (~0.15, hV4-IPS versus ~0.25, V1-V3ab). The time constant τϵis similar to that observed for the normalization fit. *(D) Summary metrics*. R_double_ is below 1 for all ROIs, indicating sub-additivity, and decreases along the visual hierarchy (V1-V3, ~0.67, LO-IPS, < 0.62). TISI is short in the earlier areas (V1-V3, ~100 ms) compared to most of the later areas. (Figure made from the script trf_mkFigure7.m.)

Finally, for completeness, we fit two other variants of the CTS model, one in which the exponent of 2 in the numerator and denominator was replaced by a free parameter, *n*,

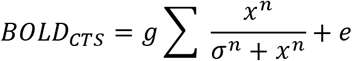

and one in which the exponents in the numerator and denominator were fit with separate free parameters, *m* and *n*:

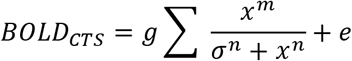

These two variants of the CTS model increase the number of free parameters from 3 (τ, σ, and g) to 4 and 5, respectively (Figure 12A). For the implementations with more free parameters, the multiple nonlinear parameters (σϵand exponents) appear to trade-off to a certain degree, so that the error bars on the separate parameters tend to be larger than those for the simpler implementation of the CTS model. Moreover, as the models become more complex, the separate parameters are harder to interpret. For example, σϵis an order of magnitude bigger in the rightmost compared to the leftmost model, but it is also raised to a higher exponent (*n*) in the rightmost model, hence the effect of normalization may be similar for the two models (Figure 12A).

**Figure 12.**
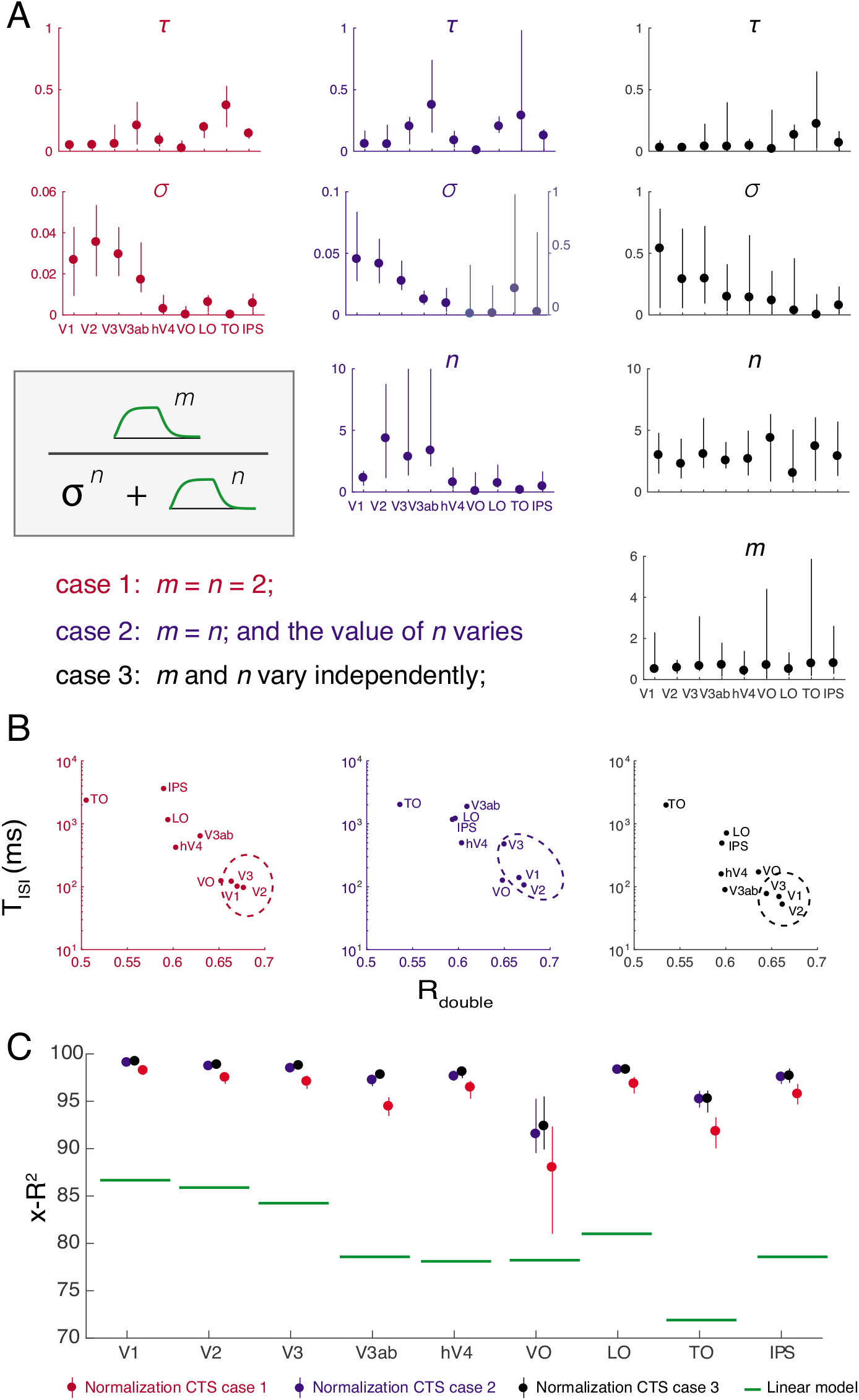
Three implementations of divisive normalization for the CTS model show the same pattern of effects. *(A)* We show three implementations of divisive normalization, with increasing numbers of free parameters from L to R: Left, the simplest implementation (same as Figures 4-10), with the exponents fixed at 2; Middle, the exponent is a free parameter; Right, the exponents in the numerator and denominator are each free parameters. In each case, the normalization step is preceded by normalization with an impulse response function and followed by scaling and summation to predict the BOLD signal, as indicated in Figure 3. Each of the three implementations is fit to the same data (Experiment 1, same as Figure 4). *(B)* The summary metrics, T_isi_ and R_double_, are similar for the three implementations, with a general tendency for V1-V3 (circled) to have shorter T_isi_ and higher R_double_, indicated by the lower right position in the scatter plots. This shows that the three CTS implementations, despite different parameterizations, manifest in similar model behavior. *(D)* Cross-validated accuracy is high for all three model forms, well above the linear model for all areas.

Because the individual model parameters are difficult to interpret, and the models differ in the number of free parameters, it is most informative to compare them on summary metrics. This shows that the same general pattern holds for all implementations: early visual areas (V1-V3) tend to have shorter Tisi and larger R_double_ (Figure 12B). Area TO is at the opposite extreme, with long T_isi_ and low R_double_. All three model variants have very high cross-validated accuracy, substantially outperforming the linear model (Figure 12C). Although the models with more parameters have numerically higher accuracy, the difference is small, and hence we tend to favor the simpler, more interpretable implementation. This bias toward simpler implementations is consistent with other uses of divisive normalization (Carandini and Heeger, 2012).

## 4. Discussion

### 4.1 Summation and adaptation in visual cortex

We report that temporal summation is subadditive throughout human visual cortex. Across 10 visual areas, BOLD responses to long stimuli were less than the linear prediction from briefer stimuli. This sub-additivity was especially pronounced in areas anterior to V1-V3. We captured this effect with a new temporal receptive field model, comprised of a linear stage followed by a static non-linearity. This compressive temporal summation model made highly accurate predictions for the fMRI data, and in all visual areas was substantially more accurate than a linear model. A single model accurately predicted two phenomena: subadditivity in the duration-response function and adaptation over short time scales (interstimulus intervals ranging from 0 to 528 ms). This indicates that both effects–the subadditivity with respect to duration and the response reduction to repeated stimuli–may arise from the same underlying processes.

A wide range of prior experimental measures are consistent with temporal subadditivities in visual cortex. For example, at the scale of 3 to 24 s, the fMRI response in V1 to a long presentation of a reversing contrast pattern is less than the prediction from a shorter presentation (Boynton et al., 1996); the fMRI response to repeated contrast patterns is larger for 1-s ISIs than 3-s ISIs (Heckman et al., 2007); the response of a V1 neuron to a steady flash is not predicted by its temporal frequency tuning and decreases over time (Tolhurst et al., 1980); the response of a neuron to a repeated stimulus is less than the response to the first stimulus (Priebe et al., 2002; Motter, 2006). Here we both quantified temporal subadditivities across the cortical visual hierarchy and account for the effects with a forward model. The model generalizes from the observed effects, as it takes arbitrary temporal patterns as inputs. The two operations – linear summation and a compressive nonlinearity – provide a simple and interpretable set of computations that can be used to characterize neural response properties across visual areas. For example, an implication of the TISI measures is that when designing an fMRI experiment, stimuli must be spaced by at least 100ms to avoid significant interactions in V1 responses, and at least 1 s in TO or IPS.

### 4.2 Subadditivities in fMRI

In principle, the subadditivity could arise from the neuronal responses, coupling between neuronal processes and the BOLD signal, or a combination of both. There are several reasons to believe that at least a significant part of the observed non-linearity is neuronal in origin. First, single unit measurements of cortical neurons show temporal sub-additivities (Tolhurst et al., 1980; Motter, 2006), and it is more parsimonious to attribute subadditivities in the single unit and BOLD measurements to a single cause. Second, we find greater subadditivities in later than earlier visual areas, consistent with a cascade architecture in which later areas add additional non-linearities to the outputs from earlier areas (Heeger et al., 1996; Simoncelli and Heeger, 1998; DiCarlo et al., 2012; Kay et al., 2013a; Kay et al., 2013b); in contrast, there is no reason to expect that the coupling between neuronal signals and the hemodynamic response would become increasingly compressive along the visual hierarchy. Third, because even our longest stimuli were brief (≤ 528ms), thereby eliciting relatively small BOLD signals (~0.5%), it is unlikely that saturation of the BOLD signal for longer stimulus durations could explain the compressive response. For example, when similar stimuli are presented in a sequence of several images, the fMRI responses are several times larger (1-4%) (Kay et al., 2013a; Kay et al., 2013b), indicating that the BOLD signal measured here was well below saturation. Therefore, overall our results indicate that the neuronal response underlying the BOLD signal shows significant temporal subadditivities, and that the subadditivity is more pronounced in later visual areas.

Multiple studies are consistent with the possibility that the linear approximation of the neural-to-BOLD transform is reasonably good (Boynton et al., 1989; Heeger et al., 2000; Rees et al., 2000). However, our interpretation of temporal compressive summation in the neural response does not rely on the assumption that the BOLD signal is *exactly* a linear transform of local neuronal activity. If, for example, the coupling reflects an approximately square root compression, as recently suggested by one group (Bao et al., 2015), then the stimulus-to-BOLD nonlinearity we observed would still imply a highly compressive neural response. This is easiest to appreciate for the power-law implementation of the CTS model. For example, the median exponent fit to the BOLD signal across ROIs ranged from 0.1 (IPS) to 0.28 (V1). If we assume that this includes a neurovascular compressive exponent of 0.5, then the stimulus-to-neural response would have exponents ranging from 0.2 (IPS) to 0.56 (V1), still highly compressive. This interpretation is supported by preliminary analyses of intracranial data, which show substantial temporal non-linearities in the neural response (Zhou et al., 2017).

### 4.3 Spatial and temporal subadditivities

Subadditive temporal summation is likely to have important functional consequences. The two ways we documented temporal subadditivities, a compressive function of duration for single stimuli, and a reduced response for paired stimuli with short ISIs, are consistent with neural adaptation: a reduced response to prolonged or repeated stimuli. These phenomena are thought to reflect adaptive changes to the local environment, rather than being a passive by-product of neural responses (Webster, 2015). For example, adaptation may serve to prioritize new information or act as gain control (Solomon and Kohn, 2014). An interesting consequence of subadditive temporal summation is that responses to stimuli of different durations are more similar to one another than they would be if summation were linear. This may be thought of as a form of *duration or timing tolerance*, analogous to size and position tolerance in spatial encoding, which are increasingly prominent in higher visual areas (Kay et al., 2013a). For example, in V1, as the stimulus size increases or the stimulus duration lengthens, the response amplitude increases substantially, whereas in area TO, the response amplitudes increase only slightly, indicating greater tolerance for size and duration (Figure 13).

**Figure 13.**
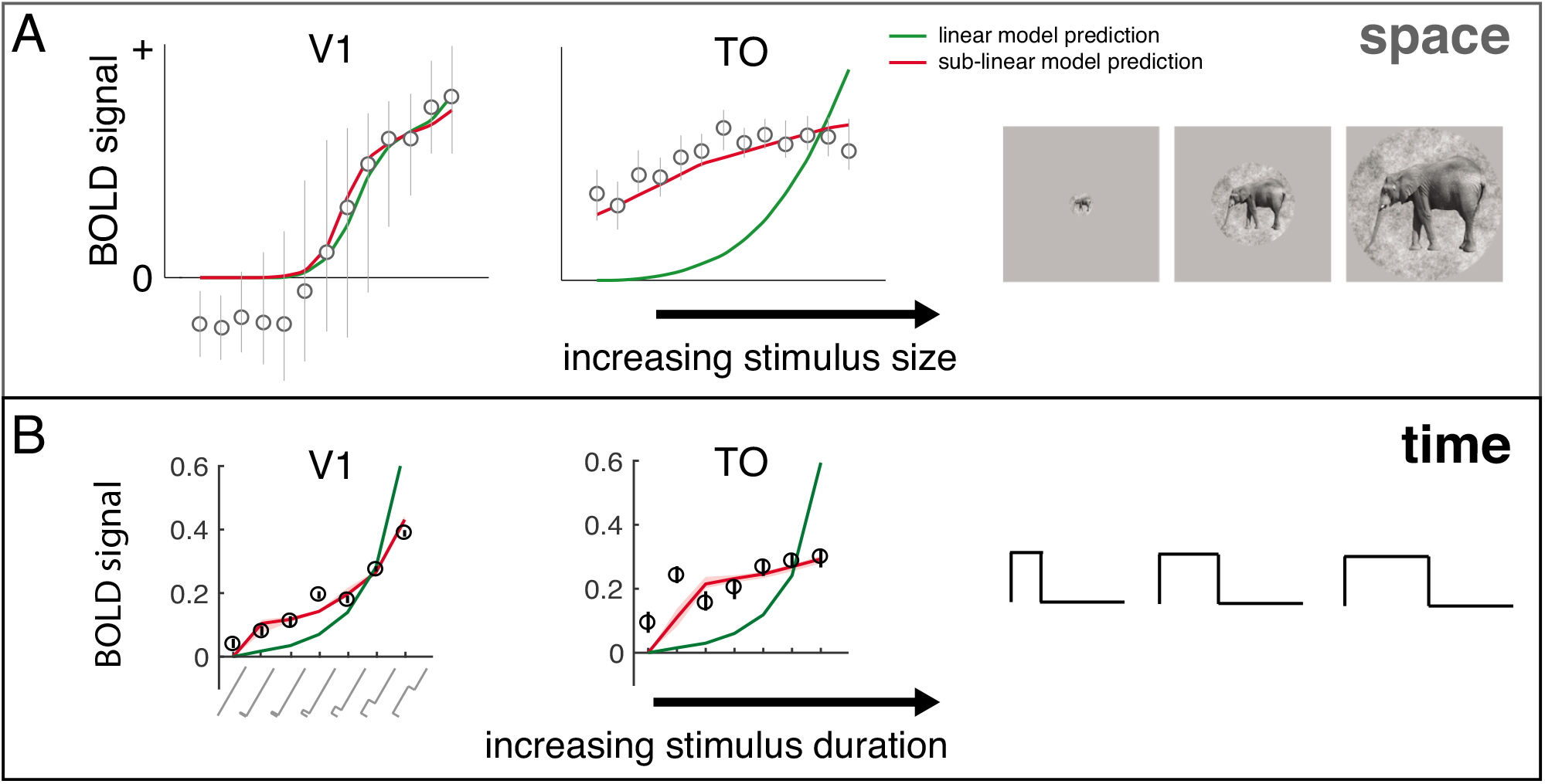
Sub-additive spatial and temporal summation. (A) BOLD responses pooled across voxels in V1 (left) and in TO (right) are plotted as a function of stimulus size. Circles and error bars are means and standard errors across bootstrapped estimates. A compressive spatial summation model (red), fit to separate data, predicts the responses slightly more accurately than a linear model (green) in V1, and substantially more accurately in TO. Adapted from figure 8 in (Kay et al., 2013a). (B) A similar pattern is observed for duration, replotted from Figure 4A.

While spatial and temporal subadditivities share some properties, they are independent findings and differ in detail. For example, V2 shows substantially more spatial subadditivity than V1 (fig 9b in (Kay et al., 2013b); fig 7b in (Kay et al., 2013a)), but a similar degree of temporal subadditivity (Figures 6 and 7). Moreover, temporal subadditivities are directional: the future cannot affect the past, whereas responses to two spatial locations can affect each other. Further, a system which is space-time separable could, in principle, exhibit saturation with space but be linear in time, or vice versa. It will be important in future work to develop an integrated model which accounts for spatial and temporal nonlinearities.

### 4.4 Temporal window length

Our finding that time scales lengthen across the visual hierarchy is consistent with measurements of temporal dynamics at a larger scale. For example, temporal receptive window length was studied by measuring response reliability to scrambled movie segments (Hasson et al., 2008; Honey et al., 2012): In visual cortex, responses depended on information accumulated over ~1s, whereas in anterior temporal, parietal and frontal areas the time scale ranged from ~12-36s. Similarly, in event-related fMRI, the influence of prior trials was modeled with an exponential decay, with longer time constants in later areas: Boynton et al (1996) reported a time constant of ~1s in V1 for contrast reversing checkerboards, and Mattar et al (2016), using static face images, reported short time constants in V1 (~0.6s) and much longer constants in face areas (~5s). In macaque, the timescale of autocorrelations in spike counts was longer for areas higher in the hierarchy (~300ms) compared to sensory areas (~ 75-100ms; Murray et al., 2014). These studies used very different methods and resulted in a wide range of time-scale estimates. It will be important in future work to ask whether a forward model can account for the range of values.

Analyzing visual information at multiple temporal scales has benefits. First, accumulating information in the past is necessary for predicting the future, and a hierarchy of temporal windows may be useful for predictions over different time-scales (Heeger, 2017). Second, signal-to-noise ratios are optimized when the temporal scale of analysis is matched to the temporal scale of the event of interest (a “matched filter”); different visual areas extract information about different image properties, which in turn are likely to have different temporal (or spatiotemporal) distributions in natural viewing. For example, V1 cells are highly sensitive to the spatially local orientation, contrast, and spatial frequency in an image. These properties are likely to change with even small eye movements, such that integrating over too long a time period will blur the dimensions of interest. In contrast, higher order image statistics may be stable over larger image regions and longer viewing durations, and hence an area sensitive to such properties may benefit from longer periods of integration. Whether or not the time scales of the different cortical areas are fixed, or adjust based on the ongoing statistics of visual input, is an important question for future work.

Just as understanding natural image statistics may lead to better theories of neural coding (Schwartz and Simoncelli, 2001; Olshausen and Field, 1996), understanding neural coding can help us understand behavior. For example, the time-scale of cortical areas may set the time-scale of integration for behavior. Words, faces, and global motion patterns are integrated over periods 5-10 times longer than textures and local motion patterns (Holcombe, 2009). These effects have not been connected to a neural model; modeling the time-scale of cortical areas critical for these tasks may help explain these large behavioral effects.

### 4.6 Generalization and future directions

The CTS model parameters estimated from our main experiment are similar to those from the second experiment (self-replication), in which we used different stimulus images. Yet, just as with spatial pRF models, it is likely that our model will fail for certain tasks or stimuli (Wandell and Winawer, 2015). For example, sustained attention to the stimulus (Self et al., 2016), presence of a surround (Bair et al., 2003), non-separable spatiotemporal patterns (motion), and stimulus history of many seconds or more (Weiner et al., 2010), can all affect the time course, hence subadditivity of the response. By formulating a forward model of responses to large-field contrast stimuli during passive viewing, we provide a quantitative benchmark that can be used to guide interpretation of how other factors influence response dynamics, and a platform upon which to extend the model to new stimulus or task features. An important goal for future work is to develop a space-time model that simultaneously accounts for nonlinearities in spatial (Kay et al., 2013a) and temporal summation. Finally, our fMRI model contains a static nonlinearity. Measurements with finer temporal resolution such as intracranial EEG will be informative for understanding the time scale of the nonlinearities.

## Acknowledgements

We thank David Heeger, Brian Wandell, and Mike Landy for comments on an earlier draft of this manuscript. We also thank Bosco Tjan, David Heeger, XJ Wang, Denis Pelli and Rachel Denison for helpful discussions and feedback as we developed our models and analyses. The research was supported by NIH grants R00-EY022116 and R01-MH111417 (J.W.)

